# Targeting STING in experimental peritoneal damage: a novel approach in peritoneal dialysis therapy

**DOI:** 10.1101/2024.11.19.621845

**Authors:** Vanessa Marchant, Jorge García-Giménez, Guadalupe T. González-Mateo, Pilar Sandoval, Lucía Tejedor-Santamaria, Sandra Rayego-Mateos, Ricardo Ramos, José Jiménez-Heffernan, Alberto Ortiz, Anne-Catherine Raby, Manuel López-Cabrera, Adrián M. Ramos, Marta Ruiz-Ortega

## Abstract

Peritoneal dialysis (PD) is a widely used kidney replacement therapy for end-stage kidney disease (ESKD) patients. However, long-term exposure to PD fluids (PDF) can lead to peritoneal membrane (PM) damage, causing ultrafiltration failure and thus PD discontinuation. Investigating the molecular mechanisms underlying this damage is crucial for identifying new therapeutic targets to mitigate peritoneal deterioration in PD patients. Therefore, in this work we study the role of STING in peritoneal inflammation and fibrosis. To this aim, we performed different preclinical mouse models of peritoneal inflammation, fibrosis, and adhesions. In a chlorhexidine gluconate (CHX)-induced inflammation model, we found changes in the peritoneal transcriptomic profile, and cytosolic DNA-sensing signaling was one of the most enriched KEGG pathways. STING, as a conspicuous member of this pathway, was upregulated in CHX-and PDF-exposed mice, and in peritoneal biopsies from PD patients. STING genetic deficiency diminished peritoneal inflammation, by downregulating inflammatory gene expression, preventing NF-κB pathway activation, and decreasing cell infiltration, in early (10 days) and advanced (30 days) stages of the CHX model. STING absence also decreased PM thickness and fibrosis in the advanced CHX model, reduced adhesion scores in a post-surgical intra-abdominal adhesion model, and decreased inflammation in an *S. epidermidis*-induced peritonitis model. Furthermore, pharmacological inhibition of STING with C-176 decreased inflammation and macrophage-mediated mesothelial-to-mesenchymal transition in cultured mesothelial cells, and reduced CHX-induced PM thickness and inflammation in mice. Altogether, these findings highlight STING as a key mediator of peritoneal damage and suggest it may be a novel therapeutic target for preventing PD-associated peritoneal deterioration.

## Introduction

Chronic kidney disease (CKD) is a health problem whose prevalence in the population has increased considerably in recent years. Currently, more than 850 million people suffer from CKD in the world [1] and it is estimated that it will become the fifth cause of death in 2040 [2]. Current therapeutic strategies only slow the progression of CKD, but many patients still progress to end-stage kidney disease (ESKD), requiring kidney replacement therapy (KRT) and developing cardiovascular complications and other comorbidities [3,4]. Peritoneal dialysis (PD) is a KRT that takes advantage of the peritoneum as a semipermeable membrane for the exchange of solutes and water between the blood in the peritoneal capillaries and the peritoneal dialysis fluid (PDF) infused into the peritoneal cavity [5]. However, the repeated and prolonged exposure of the peritoneum to conventional PDFs, that use glucose as an osmotic agent, induces peritoneal injury that leads to ultrafiltration failure and therapy discontinuation [6]. PDF exposure induces a series of cellular and molecular changes in the peritoneal membrane (PM), including loss of mesothelial cells (MCs) by death, detachment or mesothelial-to-mesenchymal transition (MMT), and sustained inflammation, characterized by macrophages and T cells recruitment, linked to the activation of different intracellular pathways, such as nuclear factor (NF)-κB signaling [7]. These pathological responses lead to submesothelial fibrosis, intra-abdominal adhesions formation, calcification and neovascularization [8–10]. In addition, peritoneal infections, mainly bacterial, may also occur, leading to recurrent peritonitis episodes that contribute to aggravating the local and systemic damage [10]. Although the molecular mechanisms involved in these harmful processes have been studied, they are still not fully understood. Thus, studying the mechanisms of peritoneal damage and finding new mediators that allow the development of new therapeutic strategies is essential.

Activation of innate immunity is critical for the clearance of infectious agents but is also involved in the response to damage-associated molecular patterns (DAMPs) released by stressed or damaged cells [11,12]. Exogen DNA from viruses or bacteria and DNA released from stressed mitochondria and damaged genomic DNA can act as DAMPs that are recognized by cytosolic DNA sensors, such as the stimulator of interferon genes (STING) [13]. STING recognizes the second messenger cGAMP, which is synthesized by the nucleotidyltransferase cGAS after interaction with cytoplasmic dsDNA [14,15]. Following interaction with cGAMP, STING translocates from the endoplasmic reticulum to the ERGIC/Golgi, where it interacts with the kinase TBK1 and the transcription factor IRF3 to induce the synthesis and secretion of type 1 interferons (IFNs) [16,17]. Type 1 IFNs then bind to the IFNAR receptor and activate a set of interferon-stimulated genes (ISGs) [18,19]. Together with the IFN response, STING recruits the IκB kinase ε (IKKε) to activate the NF-κB pathway to induce the transcription of a series of pro-inflammatory cytokines and chemokines in immune and epithelial cells [17,20]. STING hyperactivation is involved in autoinflammatory and autoimmune diseases and is increasingly related to several chronic inflammatory conditions [21]. In the peritoneum, STING deficiency, specifically in endothelial cells, has been described to prevent T cell recruitment to the peritoneal cavity in a mouse model of TNF-α-induced peritonitis [22]. However, the role of STING in PD-associated peritoneal damage has not been studied. Here, we report the activation of STING and dependent downstream pathways in response to experimental peritoneal injury of different etiology and in peritoneal biopsies of PD patients. In addition, we show that genetic STING deficiency or its pharmacological inhibition alleviate peritoneal inflammation, fibrosis and intra-abdominal adhesions in preclinical peritoneal damage models, including sterile and infectious peritoneal injury, and peritoneal adhesions. Therefore, we propose that STING may represent a promising novel therapeutic target to prevent peritoneal deterioration associated with the complications of PD treatment in ESKD patients.

## Materials and methods

### Ethical statement

Experiments on peritoneal biopsies from patients were performed according to the Declaration of Helsinki guidelines. Written informed consent was obtained from all patients prior to samples obtention and approved by the Ethics Committee of Hospital Universitario La Paz, Madrid, Spain (HULP PI-4600; Ref. 07/253477.9/21).

All animal procedures were performed in mice according to the European Community and ARRIVE reporting guidelines for the care and use of laboratory animals, with the prior approval by the Animal Ethics Committee of the IIS-FJD and Comunidad Autónoma de Madrid (PROEX 242.2/21).

### Human samples

Formalin-fixed paraffin-embedded peritoneal biopsies were used for STING immunostaining as described below. Biopsies were obtained from different PD patients at the time of kidney transplantation, nephrectomy, or catheter replacement. Control biopsies were obtained from hemodialysis patients at the time of kidney transplantation or pre-dialysis patients at the time of catheter insertion for PD.

### Animals

C57BL/6J wild-type (WT) and C57BL/6J-STING1gt/J (systemic STING deficiency) male and female mice (8–12-week-old) were obtained from Charles River Laboratories Spain (Barcelona, Spain) and maintained at the IIS-Fundación Jiménez Díaz animal facilities, with free access to food and water, normal light/dark cycles, and under specific pathogen-free conditions.

### Chlorhexidine gluconate-induced peritoneal damage mouse model

Peritoneal damage was induced in mice by daily intraperitoneal injections of 0.1% chlorhexidine gluconate (CHX) dissolved in saline solution (0.9% NaCl) at a volume of 10 mL/kg of weight, as previously described [23]. CHX was delivered daily during different time periods, as needed. Mice were randomized by assigning them to the different experimental groups. To evaluate the progression of the CHX-induced peritoneal damage, mice were non-injected (n = 6, control group), or injected with CHX for 3, 10, or 30 days (n = 6 each group). Then, two different CHX exposure models were performed to evaluate the effect of STING genetic depletion. A 10-day-CHX exposure model was performed in WT and STING-deficient (STING-KO) male mice to evaluate inflammation (WT Control group: n = 6; WT CHX group: n = 8; STING-KO Control group: n = 4; and STING-KO CHX group: n = 4). A 30-day-CHX exposure model was performed in WT and STING-deficient (STING-KO) male mice to evaluate fibrosis (WT Control group: n = 5; WT CHX group: n = 8; STING-KO Control group: n = 5; and STING-KO CHX group: n = 9). In addition, a 10-day-CHX infusion model was performed in parallel with administration of the STING inhibitor C-176 (750 nmol/0.2 mL/mouse, dissolved in corn oil as vehicle), vehicle (alone or together with CHX), or CHX alone. The experimental groups of this model were: Control (n = 6), Vehicle (n = 4), CHX (n = 6); CHX + Vehicle (n = 6), and CHX + C-176 (n = 7). Mice from all CHX models were euthanized, peritoneal lavages were obtained, and parietal peritoneal tissue samples were collected according to the specific requirements for the subsequent analyses.

### Post-surgical intra-abdominal adhesions mouse model

A standard surgical protocol was employed to perform the peritoneal adhesion mouse model, as previously described [24]. Briefly, 8-10-week-old WT (n = 5) and STING-KO (n = 6) male mice were anaesthetized with inhaled isoflurane during surgery. WT (n = 5) and STING-KO (n = 6) mice without surgery were used as controls. After accessing the abdominal cavity, three ischemic buttons (IBs) were made on the left side of the peritoneum by taking 3 mm of peritoneal parietal tissue with a hemostatic forceps and ligating the base of each segment with 4–0 silk sutures. In addition, the cecum was isolated and gently rubbed with sterile cotton swabs to induce injury in the visceral peritoneum and promote adhesions. After the procedure, the incision was closed, and mouse recovery was monitored. Mice were euthanized 5 days after surgery and the adhesions formed were quantified and scored as formerly described [24]. The extent (grade) and the quality (tenacity) of adhesions were scored for each IB with a point scale (Table 1). After examination, the parietal peritoneal tissue (IB and non-IB tissue) samples were collected according to the specific requirements for the subsequent analyses.

**Table 1.** Post-surgical adhesion mouse scoring scheme.

| Grade | Tenacity |
| --- | --- |
| (0) 0% | (0) No adhesion |
| (1) < 25% | (1) Adhesion fell apart |
| (2) 25–50% | (2) Adhesion lysed with traction |
| (3) 51–75% | (3) Adhesion lysed with blunt dissection |
| (4) >75% | (4) Adhesion lysed with sharp dissection |
|  | (5) Adhesion between the abdominal wall and organs |

### *S. epidermidis*-induced peritonitis mouse model

To induce bacterial peritonitis, 8-10-week-old WT (n = 6) and STING-KO (n = 4) female mice were intraperitoneally injected with a single dose of live *Staphylococcus epidermidis* (5 x 10^8^ cfu/mouse), as previously described [25,26]. WT (n = 6) and STING-KO (n = 6) mice without treatment were used as controls. Live *S. epidermidis* inoculum was prepared based on previously established procedures [27]. 72 hours after bacterial injection, mice were euthanized, and parietal peritoneal tissue samples were collected according to the specific requirements for the subsequent analyses.

### Transcriptomic analyses

The RNA-seq study was carried out starting from peritoneal tissue samples of C57BL6/J male mice intraperitoneally injected with 0.1% CHX daily for 10 days (n = 4) and non-injected mice as control (n = 4). RNA extraction, library preparation, and sequencing were performed in Fundación Parque Científico de Madrid (FPCM). Briefly, mechanical disruption in a TissueLyser homogenizer (Qiagen; Hilden, Germany) and RNeasy Mini Kit (Qiagen; Hilden, Germany), including on-column DNase treatment, were used for total RNA extraction from parietal peritoneal tissues, following manufacturer’s recommendations. Total RNA quality and concentration were assessed on 4200 TapeStation (Agilent Technologies; Santa Clara, CA, USA) using an RNA ScreenTape, confirming all samples had RIN > 7. Then, 500 ng of total RNA from each sample were used as input for library preparation with NEBNext Ultra II Directional RNA Library Prep Kit (New England Biolabs; Ipswich, MA, USA), following manufacturer’s recommendations for Poly(A) mRNA protocol. The finally obtained libraries were validated and quantified by TapeStation and an equimolecular pool was made, purified using AMPure XP beads (Beckman Coulter; Brea, CA, USA) and titrated by PicoGreen. The library pool was sequenced on a NextSeq P4 flowcell (Illumina; San Diego, CA, USA) where clusters were formed and sequenced in a 1x75 single-read sequencing run on a NextSeq 2000 sequencer (Illumina; San Diego, CA, USA). Read cleaning was performed with PrinSeq. Mapping and alignments to mouse genome were made with TopHat (v2.1.1). Differential expression analysis and statistics were carried out with CuffDiff from Cufflinks.

### Functional enrichment analyses

Differentially expressed genes (DEGs) found in CHX-exposed mice versus control mice (obtained from the transcriptomic analysis) were used to perform a series of functional enrichment analyses using different web tools and databases. Principal component analysis (PCA), heatmapping, and hierarchical clustering of DEGs were carried out using the ClustVis web tool (https://biit.cs.ut.ee/clustvis/) [28]. For principal component analysis (PCA), unit variance scaling is applied to rows; singular value decomposition with imputation is used to calculate principal components. For hierarchical clustering, rows were centered and unit variance scaling was applied to rows. Both rows and columns were clustered using correlation distance and average linkage. Functional enrichment analysis of cell type and transcription factors was performed through the Metascape web tool (https://metascape.org/) [29], based on the Pattern Gene Database (PaGenBase) and Transcriptional Regulatory Relationships Unraveled by Sentence-based Text mining (TRRUST) databases. Signaling pathway enrichment analysis was carried out using the g:Profiler web server (https://biit.cs.ut.ee/gprofiler/gost) [30] based on the Reactome and Kyoto Encyclopedia of Genes and Genomes (KEGG) databases. For KEGG enrichment, terms were filtered by including the word “pathway”. Identification of upregulated DEGs on the KEEG cytosolic DNA-sensing pathway (mmu04623) was performed using the ShinyGo 0.80 web application (https://bioinformatics.sdstate.edu/go80/) [31]. All functional enrichment analyses were performed for up and downregulated DEGs separately.

### Cell Culture

The human mesothelial cell line MeT-5A (ATCC; Rockville, MD, USA) was cultured in Earle’s M199 medium supplemented with 10% FBS (Gibco; Waltham, MA, USA), 20 mM HEPES (Gibco; Waltham, MA, USA), and 100 U/mL penicillin and 100 μg/mL streptomycin (Gibco; Waltham, MA, USA). Cells were grown at 37°C, in a humidified atmosphere with 5% CO2. For *in vitro* assessment of mesothelial inflammation and MMT, MeT-5A cells were FBS-depleted for 24 hours, pre-treated with C-176 (1 μM) for 1 hour when corresponded, and then stimulated with TNF-α (5 ng/ml) or TGF-β1 (3 ng/ml) for 48 hours.

To obtain primary murine peritoneal macrophages, C57BL6/J mice were intraperitoneally injected with 3% thioglycolate to induce macrophage recruitment into the peritoneal cavity. 72 hours after, macrophages were removed by washing the peritoneal cavity with saline solution. Macrophages were cultured in RPMI 1640 medium (Gibco; Waltham, MA, USA) supplemented with 10% FBS (Gibco; Waltham, MA, USA), 2 mM L-glutamine (Euroclone; Pero, Italy), 100 U/mL penicillin, and 100 μg/mL streptomycin (Gibco; Waltham, MA, USA), and grown at 37°C, in a humidified atmosphere with 5% CO2. For the experiments, macrophages were FBS-depleted for 24 hours, pre-treated with C-176 (1 μM) for 1 hour when corresponded, and then stimulated with LPS (µg/mL) for 6 hours. After this time, cultured medium was renewed and, 24 hours later, macrophage-conditioned medium (MCM) was obtained and stored. Then, cultured MeT-5A cells (FBS-depleted for 24 h) were incubated for 48 h in the presence of MCM (from macrophages with or without LPS) and MeT-5A medium (M199), in a 1:1 proportion. MeT-5A cells incubated with RPMI medium, instead of MCM, were used as control.

### Histological and immunohistochemical analyses

Peritoneal tissues collected from mice and patients (biopsies) were fixed in 4% formaldehyde, embedded in paraffin, and cut in 4 μm tissue sections for Masson’s trichrome staining, (Bio-Optica; Milano, Italy), following the manufacturer’s instructions, immunohistochemistry (IHC), or immunofluorescence (IF) studies.

For IHC, tissue sections were deparaffined and hydrated, and antigen retrieval was carried out using a sodium citrate buffer (10 mM, pH 6 or 9) on a PT Link system (DAKO, CA, USA). Endogenous peroxidase blockade was done using 3% hydrogen peroxide (Millipore, MA, USA) and protein blockade using a Casein blocking solution (Vector Laboratories, CA, USA). Then, primary antibodies were incubated overnight at 4°C, followed by incubation with biotinylated secondary antibodies (anti-rabbit or anti-rat, 1:200) and avidin-biotin complex (ABC, Vector Laboratories; CA, USA). Signal was detected using 3,3-diaminobenzidine (DAB) chromogen and substrate solution (Abcam; Cambridge, UK). Finally, slides were counterstained with hematoxylin (Merck; Darmstadt, Alemania), dehydrated and mounted with DPX (Merck; Darmstadt, Alemania). The following primary antibodies were used: rabbit anti-STING (1:500; #13647, Cell Signaling; Danvers, MA, USA), rat anti-F4/80 (1:50; MCA497, Bio-Rad; CA, USA), mouse anti-α-SMA (1:200; A2527, Sigma-Aldrich; Saint Louis, MO, USA), mouse anti-phospho(S536)-NF-κB p65 (1:50; sc-136548, Santa Cruz Biotechnology; Dalla, TX, USA), rabbit anti-MPO (1:3000; A0398, Dako; CA, USA), rabbit anti-CD3 (1:100; A0452, Dako; CA, USA), mouse anti-CD4 (1:100; M7310, Dako; CA, USA), and rabbit anti-phospho(S40)-NRF2 (1:2000; AB76026, Abcam; Cambridge, UK).

For IF, deparaffination, hydration, and antigen retrieval of tissue sections were done as above described. Next, tissues were permeabilized with 0.2% Triton X-100/PBS and protein blockade was done with 10% rabbit or rat serum, as appropriate (diluted in 4% BSA in PBS). Then, sections were incubated overnight at 4°C with the primary antibodies diluted in 1% serum (diluted in 4% BSA in PBS) followed by incubation with the fluorophore-conjugated secondary antibodies. The primary antibodies used were rabbit anti-STING (1:200; PA5-23381, Invitrogen; Waltham, MA, USA) and rat anti-F4/80 (1:50; MCA497, Bio-Rad; CA, USA), and the secondary antibodies were Alexa Fluor 633-conjugated anti-rabbit (1:200; Thermo Fisher Scientific; Waltham, MA, USA) and Alexa Fluor 488-conjugated anti-rat (1:200; A21208, Thermo Fisher Scientific; Waltham, MA, USA). Nuclei were stained with 4’,6-diamidino-2-phenylindole (DAPI, 1:10,000; Sigma-Aldrich; Saint Louis, MO, USA) and then, stained sections were mounted with ProLong Gold antifade reagent (Invitrogen; Waltham, MA, USA) and images were acquired using fluorescence microscope (BX53, Olympus; Tokyo, Japan).

Peritoneal membrane thickness was determined by measuring the submesothelial zone width on Masson stain images using ImageJ tool (five measurements/field in 5-10 fields per mouse, 200 × magnification) and IHC quantification was made by counting positive-staining in 5–10 randomly chosen fields per mouse (200 × magnification).

### Flow cytometry analysis

Cell suspensions obtained from peritoneal lavages were counted with a Scepter handheld automatic cell counter (Millipore; Burlington, MA, USA). Then, 1 x 10^6^ cells were stained with fluorochrome-conjugated mouse-specific antibodies against CD3, CD4, CD8, CD11b, F4/80, and Ly6G (BD Biosciences Pharmingen, San Diego, CA, USA) following manufacturer’s instructions. An isotype control was used in each case to ensure staining specificity and avoid artefacts. Unstained cells were used as negative control for the analysis settings. Samples were analyzed in a BD FACS Canto II (BD Biosciences, San Jose, CA) flow cytometer, and data analyses were performed using FlowJo software (Version 10).

### Protein level studies

Total protein from frozen peritoneal tissue and cultured cells were isolated by homogenization in a lysis buffer (T-PER, Thermo Fisher Scientific; Waltham, MA, USA) with 10 μL/mL protease inhibitor cocktail, 0.2 mmol/L phenylmethylsulfonyl fluoride, and 0.2 mmol/L orthovanadate. Proteins were quantified using a Pierce BCA protein assay kit (Thermo Fisher Scientific; Waltham, MA, USA) and then were separated by electrophoresis using 8–10% polyacrylamide-SDS gels under reducing conditions for western blotting. Samples were then transferred onto polyvinylidene difluoride membranes (Thermo Fisher Scientific; Waltham, MA, USA), blocked with 5% non-fat milk, and incubated overnight at 4°C with the following primary antibodies: rabbit anti-STING (1:1000; #13647, Cell Signaling; Danvers, MA, USA), rabbit anti-TBK1 (1:1000; #38066, Cell Signaling; Danvers, MA, USA), rabbit anti-phospho(S172)-TBK1 (1:1000; #5483, Cell Signaling; Danvers, MA, USA), rabbit anti-IRF3 (1:1000; #4302, Cell Signaling; Danvers, MA, USA), rabbit anti-phospho(S396)-IRF3 (1:1000; SAB4504031, Sigma-Aldrich; Saint Louis, MO, USA), rabbit anti-phospho(S536)-NF-κB p65 (1:1000; #3031, Cell Signaling; MA, USA), mouse anti-phospho(S32)-IκBα (1:1000; sc-8404, Santa Cruz; CA, USA), and rabbit anti-Fibronectin (1:5000; AB2033, Millipore; MA, USA). Then, the membranes were incubated with HRP-conjugated secondary antibodies (anti-rabbit or anti-mouse, 1:5000). Loading controls were performed using a mouse anti-GAPDH antibody (1:5000; CB1001, Millipore; MA, USA) and mouse α-Tubulin (1:5000; T5168, Sigma-Aldrich; Saint Louis, MO, USA). Proteins on membranes were visualized using the chemiluminescence detection kit Immobilon Crescendo Western HRP substrate (Millipore; Burlington, MA, USA) on the Amersham Imager 600 instrument (GE Healthcare; Chicago, IL, USA). Images were analyzed by densitometry using the image processing program written on Java, ImageJ.

### Gene Expression assays

Total RNA was isolated from frozen peritoneal tissue and pelleted cultured cells by homogenization with TRItidy G (PanReac; Darmstadt, Germany), according to the manufacturer’s protocols. Next, cDNA was synthesized using a High-Capacity cDNA Reverse Transcription Kit (Applied Biosystems; Waltham, MA, USA) starting from 2 µg of total RNA. Gene expression analysis was determined by quantitative PCR (qPCR) using the commercial master mix Premix Ex Taq (Takara; Otsu, Japan) and predesigned TaqMan-based qPCR assays for target and housekeeping genes (Table 2) and run on the 7500 Fast Real-Time PCR System and QuantStudio 3 thermocyclers (Applied Biosystems; Waltham, CA, USA). Relative expression values were normalized by the expression of housekeeping genes using the 2^-ΔΔCt^ method. The results were expressed as fold change (n-fold) relative to control.

**Table 2.** Pre-designed assays used for qPCR.

| **Specie** | **Gene** | **Gene type** | **Assay ID** | **Company** |
| --- | --- | --- | --- | --- |
| Mouse | *Acta2* | Target | Mm.PT.58.16320644 | Integrated DNA Technologies |
| Mouse | *Arg1* | Target | Mm.PT.58.8651372 | Integrated DNA Technologies |
| Mouse | *Arg2* | Target | Mm.PT.58.31712462 | Integrated DNA Technologies |
| Mouse | *Cat* | Target | Mm.PT.58.12825133 | Integrated DNA Technologies |
| Mouse | *Ccl2* | Target | Mm.PT.58.42151692 | Integrated DNA Technologies |
| Mouse | *Ccl5* | Target | Mm.PT.58.43548565 | Integrated DNA Technologies |
| Mouse | *Ccl8* | Target | Mm01297183_m1 | Applied Biosystems |
| Mouse | *Ccl19* | Target | Mm00839966_g1 | Applied Biosystems |
| Mouse | *Cd163* | Target | Mm00474091_m1 | Applied Biosystems |
| Mouse | *Cdh2* | Target | Mm.PT.58.12378183 | Integrated DNA Technologies |
| Mouse | *Col1a1* | Target | Mm.PT.58.7562513 | Integrated DNA Technologies |
| Mouse | *Col1a2* | Target | Mm.PT.58.5206680 | Integrated DNA Technologies |
| Mouse | *Cxcl1* | Target | Mm04207460_m1 | Applied Biosystems |
| Mouse | *Cxcl10* | Target | Mm.PT.58.43548565 | Integrated DNA Technologies |
| Mouse | *Hmox1* | Target | Mm.PT.58.8600055 | Integrated DNA Technologies |
| Mouse | *Il1b* | Target | Mm.PT.58.41616450 | Integrated DNA Technologies |
| Mouse | *Il10* | Target | Mm01288386_m1 | Applied Biosystems |
| Mouse | *Il6* | Target | Mm.PT.58.10005566 | Integrated DNA Technologies |
| Mouse | *Ifi44* | Target | Mm.PT.58.12162024 | Integrated DNA Technologies |
| Mouse | *Ifit1* | Target | Mm.PT.58.32674307 | Integrated DNA Technologies |
| Mouse | *Ifna1* | Target | Mm.PT.58.43426930.g | Integrated DNA Technologies |
| Mouse | *Ifnb1* | Target | Mm.PT.58.30132453.g | Integrated DNA Technologies |
| Mouse | *Ifng* | Target | Mm.PT.58.41769240 | Integrated DNA Technologies |
| Mouse | *Snai1* | Target | Mm00441533_g1 | Applied Biosystems |
| Mouse | *Mx2* | Target | Mm.PT.58.29837402 | Integrated DNA Technologies |
| Mouse | *Nfe2l2* | Target | Mm.PT.58.29108649 | Integrated DNA Technologies |
| Mouse | *Nox1* | Target | Mm.PT.58.29694286 | Integrated DNA Technologies |
| Mouse | *Nox4* | Target | Mm.PT.58.8820983 | Integrated DNA Technologies |
| Mouse | *Oasl2* | Target | Mm01201449_m1 | Integrated DNA Technologies |
| Mouse | *Sod1* | Target | Mm.PT.58.12368303 | Integrated DNA Technologies |
| Mouse | *Sod2* | Target | Mm.PT.58.14276358 | Integrated DNA Technologies |
| Mouse | *Sting1* | Target | Mm.PT.58.12798185 | Integrated DNA Technologies |
| Mouse | *Tgfb1* | Target | Mm.PT.58.11254750 | Integrated DNA Technologies |
| Mouse | *Usp18* | Target | Mm.PT.58.28965870 | Integrated DNA Technologies |
| Mouse | *Ppia* | Housekeeping | Mm.PT.39a.2.gs | Integrated DNA Technologies |
| Mouse | *Gapdh* | Housekeeping | Mm.PT.39a.1 | Integrated DNA Technologies |
| Human | *CCL2* | Target | Hs.PT.58.45467977 | Integrated DNA Technologies |
| Human | *CCL5* | Target | Hs.PT.58.1724551 | Integrated DNA Technologies |
| Human | *CDH2* | Target | Hs.PT.58.26024443 | Integrated DNA Technologies |
| Human | *CXCL10* | Target | Hs.PT.58.3790956.g | Integrated DNA Technologies |
| Human | *FN1* | Target | Hs.PT.58.40005963 | Integrated DNA Technologies |
| Human | *GREM1* | Target | Hs.PT.58.21084086 | Integrated DNA Technologies |
| Human | *SNAI1* | Target | Hs.PT.58.2984401 | Integrated DNA Technologies |
| Human | *TGFB1* | Target | Hs.PT.58.39813975 | Integrated DNA Technologies |
| Human | *USP18* | Target | Hs.PT.58.25205207 | Integrated DNA Technologies |
| Human | *GAPDH* | Housekeeping | Hs.PT.39a.22214836 | Integrated DNA Technologies |

### Statistical Analysis

In most of the figures the results are expressed as fold change (n-fold) relative to the control and represented as mean ± SEM. The Shapiro–Wilk test was used to evaluate sample normality distribution. To assess statistical differences between groups, several tests were used based on data characteristics. For two-group comparisons, F-test was used to check for homogeneity of variances; unpaired T-tests were conducted for normal samples (with or without Welch’s correction depending on variance); and Mann-Whitney test was used for non-normal samples. For more than two groups, Bartlett’s test assessed homoscedasticity; parametric one-way ANOVA followed by Fisher’s LSD test was used for normal homoscedastic samples; for heteroscedastic samples, Welch-corrected one-way ANOVA and Brown-Forsythe tests were followed by Welch’s T-tests; non-parametric Kruskal-Wallis test was applied for non-normal samples, followed by Dunn’s test; and two-way ANOVA followed by Fisher’s LSD was used for two factors (treatment and genotype). For cell culture experiments, paired comparisons were done using RM one-way ANOVA for normal data or Friedman test for non-normal data, followed by Fisher’s LSD or Dunn’s test, respectively. All statistical analyses and graphs were performed using GraphPad Prism 8.0 (GraphPad Software, San Diego, CA, USA). *P*-values < 0.05 were considered statistically significant.

## Results

### Evolution of CHX-induced peritoneal damage in mice

Chlorhexidine gluconate (CHX) is a commonly utilized agent for the experimental induction of peritoneal fibrosis in murine models, facilitating the investigation of diverse mechanisms underlying tissue damage [32]. In our study, we evaluate the time-course peritoneal injury caused by daily intraperitoneal injections of 0.1% CHX for 3, 10, and 30 days in C57BL6/J mice. The histological analysis revealed that as early as three days of CHX exposure, there was a modest but statistically significant increase in the thickness of the submesothelial zone (hereafter referred to as PM thickness). This increase was more pronounced after 10 days and further intensified after 30 days, as seen in Masson’s trichrome stained tissue sections (Supplementary Figure S1A). At the molecular level, the expression of genes associated with the MMT process and fibrosis exhibited a gradual increase in response to prolonged CHX exposure in the peritoneal tissue of mice (Supplementary Figure S1B). Accordingly, at 3 days, exposure to CHX significantly increased transforming growth factor β (TGF-β, *Tgfb1* gene) levels, whereas other markers of MMT, such as Snail and N-cadherin (*Snai1* and *Cdh2* genes, respectively), and fibrosis, like type I collagen (*Col1a1*), were not modified. After 10 days of CHX exposure, the aforementioned genes underwent upregulation, with even higher expression levels after 30 days. Regarding inflammatory markers, gene overexpression of the chemokine CCL5 (*Ccl5*) and the cytokines IL-1β (*Il1b*) and IL-6 (*Il6)* was observed as early as 3 days of CHX administration, with a sustained elevation observed at 10 days and a subsequent decline (Supplementary Figure S1C). Taken together, these results show that CHX exposure causes significant histological and molecular alterations in the mouse peritoneum, detectable as early as 3 days. The top of inflammation appears to occur after 10 days of CHX exposure, while the development of fibrosis is exacerbated by chronic exposure for 30 days. Therefore, a 10-day CHX exposure period was selected as an equitable model for the investigation of the mechanisms underlying both inflammation and early fibrosis in peritoneal damage.

### Transcriptomic changes during CHX-induced inflammation in mice

To profile the transcriptomic changes in CHX-induced peritoneal injury and help to identify new mediators of damage, we performed an RNA-seq study in the parietal peritoneum from 10-day CHX-exposed and control mice. From a total of 12.058 detected genes (Figure 1A), we found 1.050 differentially expressed genes (DEGs), 974 of them upregulated and 76 downregulated (Log_2_(FC)| ≥ 2 and *q*-value < 0.05) (Figure 1B). Principal component analysis (PCA) of DEGs, accurately clustered CHX and control groups (Figure 1C). Proinflammatory chemokines (*Ccl2* and *Ccl7*), cytokines (*Il1b*) and ISGs (*Cxcl10* and *Isg15*) highlighted among the most upregulated genes (Figure 1A). Later, a functional enrichment analysis using bioinformatic web tools and databases was conducted to explore the potential pathophysiological associations with the transcriptomic changes, particularly of upregulated DEGs. In the Metascape-linked Pattern Gene Database (PaGenBase), immune cell terms were among the most represented cell types, with macrophages being the most enriched cell subset (Figure 1D), According to the Transcriptional Regulatory Relationships Unraveled by Sentence-based Text mining (TRRUST) tool, NF-κB, STAT1, and IRF1, which are primary transcriptional commanders of the inflammatory reaction, are also predicted to be the main potential transcription factors regulating the increased gene expression in the peritoneum of CHX-exposed mice (Figure 1E). Moreover, the bulk of the most enriched Reactome pathways was also related to the immune system, particularly to the innate immune system, and associated terms of which several belong to the IFN pathway (Figure 1F). Consistent with the previous analysis, functional analysis based on the Kyoto Encyclopedia of Genes and Genomes (KEGG) database showed classical innate immune-related signaling pathways such as NOD-like receptor (NLR), Toll-like receptor (TLR), Cytosolic DNA-sensing pathway, NF-κB, and TNF-α as the most enriched terms. (Figure 1H). Altogether, these data corroborate the increased inflammatory status induced by CHX in the damaged peritoneum of mice.

**Figure 1.**
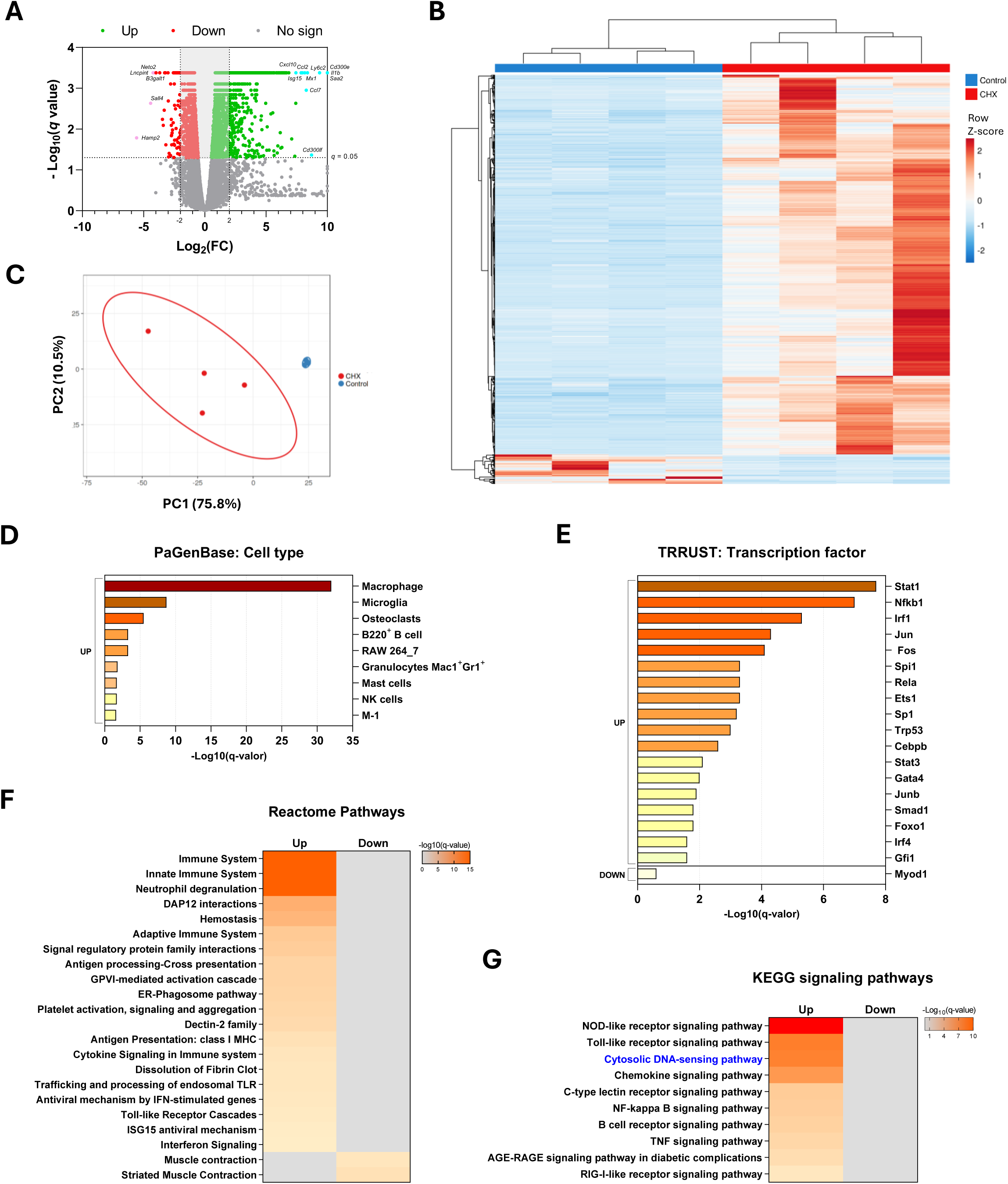
Transcriptomic and functional enrichment analysis in the peritoneum of mice exposed to CHX for 10 days. Sequencing analysis was performed starting from total RNA from parietal peritoneal tissue of male C57BL6/J mice injected intraperitoneally with 0.1% CHX for 10 days and control mice (n = 4 per group). **(A)** The volcano plot shows 12,058 genes with satisfactory statistical analysis. Differentially expressed genes (DEGs) in the CHX group versus the control group with a *q*-value < 0.05 are shown in green (increased) and red (decreased). The highest upregulated or downregulated DEGs are shown in cyan or pink, respectively. **(B)** Heatmap representing row *Z*-score and the hierarchical clustering of the 1,050 DEGs with *q*-value < 0.05 and |Log2(FC)| ≥ 2. **(C)** Principal component analysis (PCA) shows clustering within individual prediction ellipses for each group. PCA plot and heatmap were generated using the ClustVis web tool. **(D-G)** Functional enrichment analysis of cell type **(D),** transcription factors **(E)**, and signaling pathways **(F, G)** in upregulated DEGs according to Pattern Gene Database (PaGenBase), Transcriptional Regulatory Relationships Unraveled by Sentence-based Text mining database (TRRUST), Reactome, and Kyoto Encyclopedia of Genes and Genomes (KEGG) databases, respectively.

### Cytosolic DNA sensor STING is increased in peritoneal damage

Among the top KEGG signaling pathways found enriched in the functional analysis of CHX-induced upregulated genes (Figure 1H), we focused on the cytosolic DNA-sensing pathway, whose role in peritoneal damage development is still unraveled. One of the key components of this pathway is the cytosolic DNA sensor STING, which we found upregulated in the RNA-seq analysis (Supplementary Figure S2). Moreover, the increase in the STING gene expression was validated by qPCR in the peritoneum of 10-day CHX-exposed mice (Figure 2A), as well as its protein levels (Figure 2B). To evaluate STING in PD-associated damage, we performed STING immunostaining on peritoneal tissue sections from several chronic PDF-exposure murine models. STING-positive cells were abundantly found in the peritoneum of PDF-exposed mice alone (Figure 2C) or with kidney insufficiency (induced by 5/6 nephrectomy, Figure 2D), whereas STING-positive cells were scarcely present in the healthy peritoneum. To approach the clinical relevance of these findings, we studied the expression of STING in peritoneal biopsies of PD patients. STING was expressed in endothelium both in control cases and PD samples, but a marked positive staining was found in interstitial cells in the submesothelial zone of PD biopsies (Figure 2E). Therefore, STING is upregulated in the PDF-exposed peritoneum of both mice and PD patients, thus suggesting a potential role in sustained PD-associated pathophysiological mechanisms.

**Figure 2.**
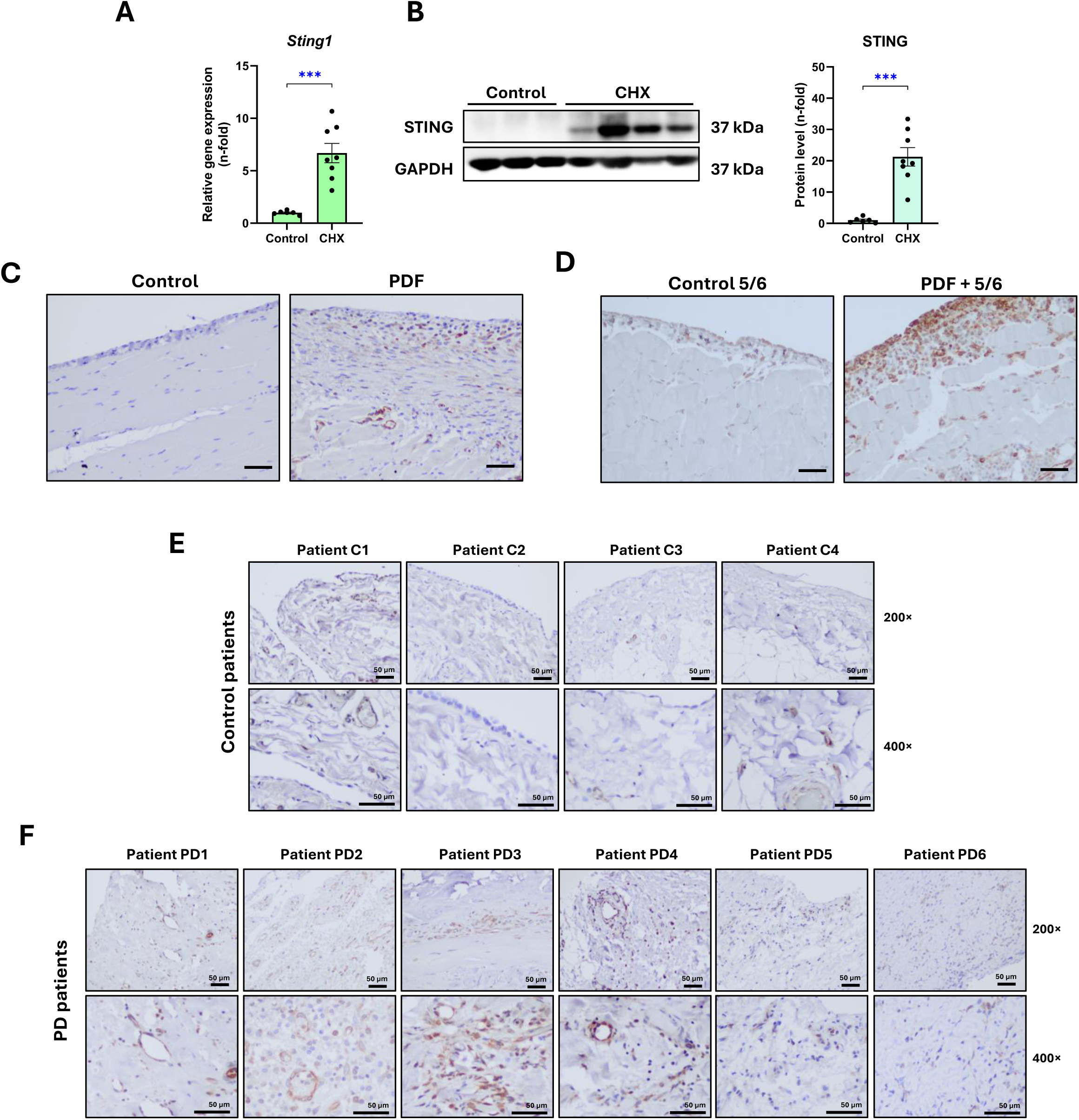
Expression of STING in the peritoneum of CHX-and PDF-exposed mice and in peritoneal biopsies from PD patients. Male C57BL/6J mice were intraperitoneally injected with 0.1% CHX for 10 days. **(A)** Relative gene expression of *Sting1* analyzed by RT-qPCR from total RNA of peritoneal parietal tissue, using *Gapdh* as housekeeping gene. **(B)** STING protein levels were assessed by western blotting from total proteins of peritoneal parietal tissue, using GAPDH as loading control. Results are expressed as fold change (n-fold) in comparison with the control group and represented as the mean ± SEM of 6-8 animals per group. *** *p* < 0.001. **(C-F)** Immunohistochemical staining of STING sections of parietal peritoneal tissue from 30-day-PDF-exposed mice **(C)**, 60-day-PDF-exposed 5/6 nephrectomized mice **(D)**, and biopsies from control **(E)** and PD **(F)** patients. Scale bar: 50 μm.

### STING deficiency ameliorates early CHX-induced peritoneal inflammation

To unravel the role of STING in the development of peritoneal damage, particularly in inflammation, we induced peritoneal injury in STING-deficient (STING-KO) mice by daily intraperitoneal exposure to CHX for 10 days. PM thickness and gene expression levels did not differ between wild-type (WT) and STING-KO control mice (Supplementary Figure S3). However, after 10-day-CHX exposure, STING-KO mice exhibited a slight but significant decrease in PM thickness compared to WT CHX mice (Supplementary Figure S3A). STING deficiency also prevented the increase in *Tgfb1* expression induced by CHX, whereas no significant effect was found in *Cdh2* and *Col1a2* gene upregulation (Supplementary Figure S3B). Regarding inflammation, macrophage tissue infiltration was lowered by STING deficiency (Figure 3A), corroborated by decreased gene expression of M1 and M2 macrophage markers, *Arg2* and *Arg1*, respectively (Figure 3B). Accordingly, STING absence fully prevented CHX-induced gene upregulation of inflammatory chemokines (*Ccl2* and *Ccl5*) and cytokines (*Il1b* and *Il6*) (Figure 3C), and ISGs (Figure 3D).

**Figure 3.**
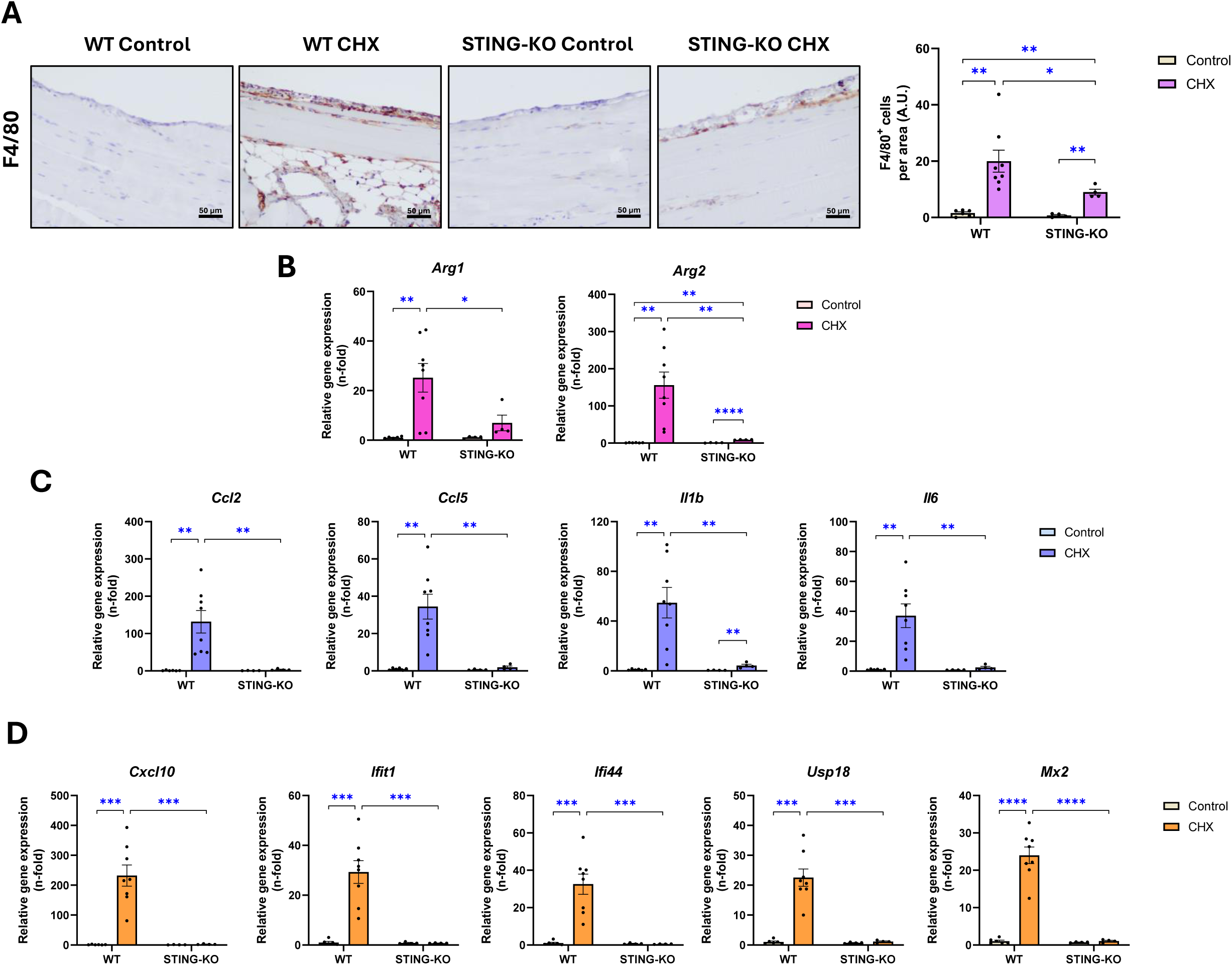
Peritoneal inflammation in WT and STING-KO mice exposed to CHX for 10 days. Male C57BL/6J wild-type (WT) and STING-deficient (STING-KO) mice were intraperitoneally injected with 0.1% chlorhexidine gluconate (CHX) for 10 days. **(A)** F4/80 immunostaining to identify macrophages infiltrates (brown-stained cells) in parietal peritoneal tissue. Microscopy images correspond to a representative animal from each group (left) and the corresponding quantification of F4/80^+^ cells (right). Scale bar: 50 μm. **(B-D)** Relative expression of inflammatory genes, including macrophage markers **(B),** inflammatory chemokines and cytokines **(C),** and interferon-stimulated genes **(D)** analyzed by RT-qPCR from total RNA of parietal peritoneal tissue, using *Gapdh* as housekeeping gene, and expressed as fold change (n-fold) relative to control. Results are represented as mean ± SEM of 4-8. * *p* < 0.05; ** *p* < 0.01; *** *p* < 0.001; and **** *p* < 0.0001. A.U.: arbitrary units.

Thus, STING deficiency prevents early peritoneal inflammation induced by CHX exposure in mice, clearly showing that STING is a key player of the inflammatory response in the peritoneum.

### STING deficiency attenuated chronic CHX-induced peritoneal fibrosis and inflammation

To better evaluate the implication ofSTING in peritoneal fibrosis and sustained inflammation, we performed an advanced 30-day CHX chronic exposure model in STING-KO. In CHX-exposed WT mice, we found abundant STING-positive cells in thickened PM and inflammatory cell infiltration areas (Figure 4A), most of them colocalizing with F4/80-positive macrophages (Figure 4B). Additionally, gene and protein STING levels were significantly increased (Figure 4C and 4D), together with increased levels of the downstream effectors of STING signaling, TBK1 and IRF3, both at basal level and in their phosphorylated and activated forms (p-TBK1 and p-IRF3) (Figure 4E and 4F). On the other hand, STING deficiency significantly reduced PM thickening induced by CHX in comparison to WT mice (Figure 5A and 5B). STING-KO mice also exhibited less peritoneal staining of the myofibroblastic cells marker, α-smooth muscle actin (α-SMA) (Figure 5C), accompanied by a decline in its gene expression (*Acta2* gene) (Figure 5D) in comparison with WT CHX mice. Moreover, the CHX-induced peritoneal overexpression of a panel of MMT markers (*Cdh2* and *Snai1*), profibrotic growth factors (*Tgfb1* and *Ccn2*) and ECM/fibrosis markers (*Col1a1*) was substantially attenuated by the STING deficiency (Figure 5E), a fact compatible with a potential STING role in promoting MMT and fibrosis development in the peritoneum. The extracellular matrix (ECM) protein fibronectin was observed to be reduced in CHX-exposed STING-KO mice in comparison to WT mice, at both the protein and gene expression levels (Figure 5F). The gene expression of the MC marker calretinin (*Calb* gene) was significantly decreased after CHX exposure in WT mice but not in STING-deficient mice (Figure 5G), suggesting that STING may play a role in MC loss.

**Figure 4.**
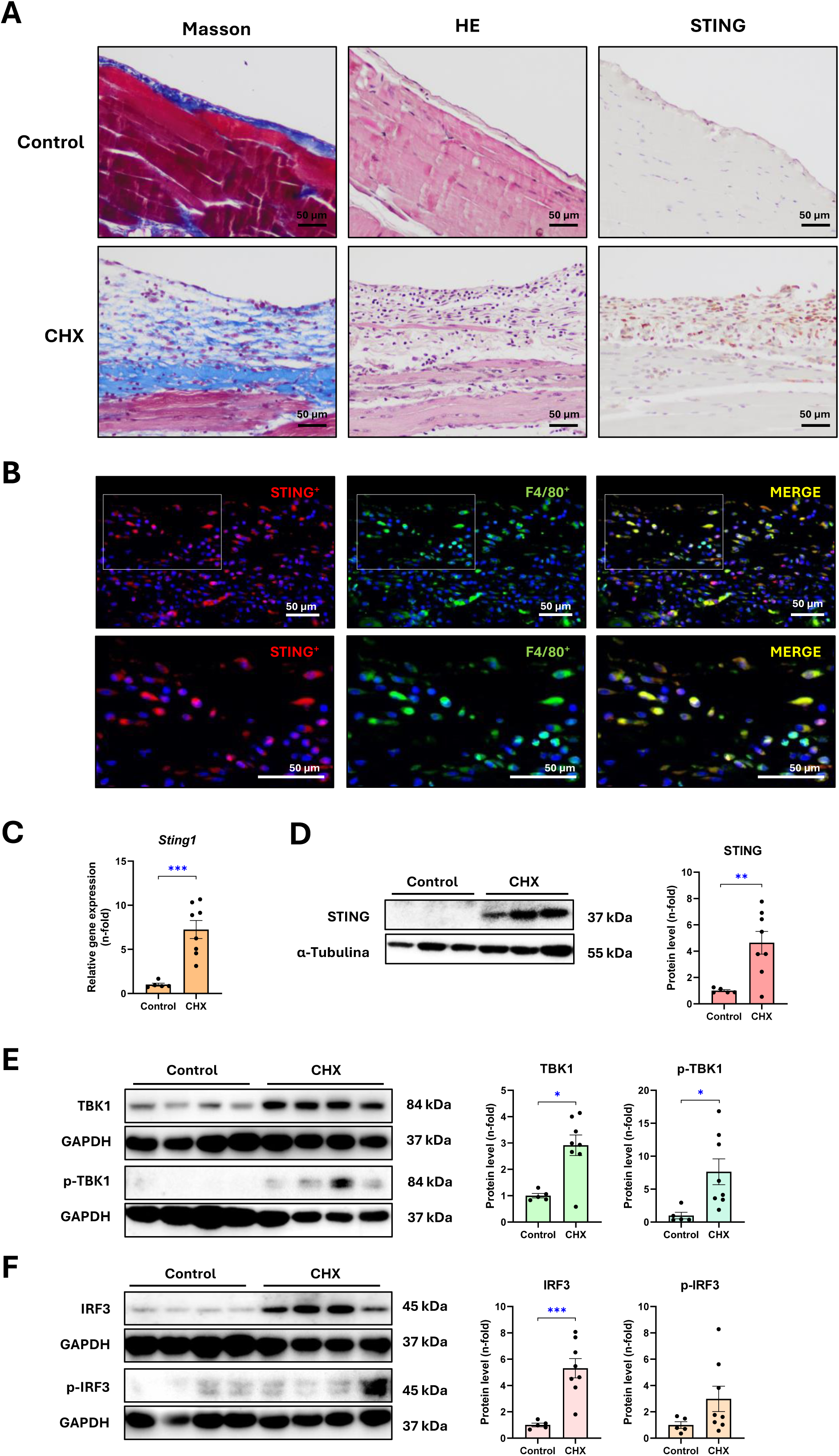
Expression and localization of STING and downstream signaling mediators in the peritoneum of mice treated with CHX for 30 days. Male C57BL/6J mice were intraperitoneally injected with 0.1% chlorhexidine gluconate (CHX) for 30 days. **(A)** Microscopy images of parietal peritoneal tissue sections stained with Masson’s trichrome and hematoxylin-eosin (HE) show thickened peritoneal membrane and increased inflammatory infiltrate in CHX-treated mice. Immunostaining shows STING^+^ cells on infiltrated and thickened peritoneal membrane of CHX-treated mice. **(B)** Immunofluorescence staining of STING (red) and F4/80 (green) shows the presence of double-labeled F4/80^+^STING^+^ (merged) macrophages in the peritoneum of CHX-exposed mice. Microscopy images correspond to a representative animal from each group. Scale bar: 50 μm. **(C,D)** Peritoneal STING gene **(C)** and protein **(D)** expression levels. **(E)** Protein levels of total TBK1 and phosphorylated TBK1 (p-TBK1). **(F)** Protein levels of total IRF3 and phosphorylated IRF3 (p-IRF3). Relative gene expression was analyzed by RT-qPCR from total RNA of parietal peritoneal tissue, using *Gapdh* as housekeeping gene. Protein levels were assessed by western blot from total proteins of parietal peritoneal tissue, using α-Tubulin or GAPDH as loading control. Results are expressed as n-fold compared to control and represented as the mean ± SEM of 5-8 animals per group. * *p* < 0.05; ** *p* < 0.01; and *** *p* < 0.001.

**Figure 5.**
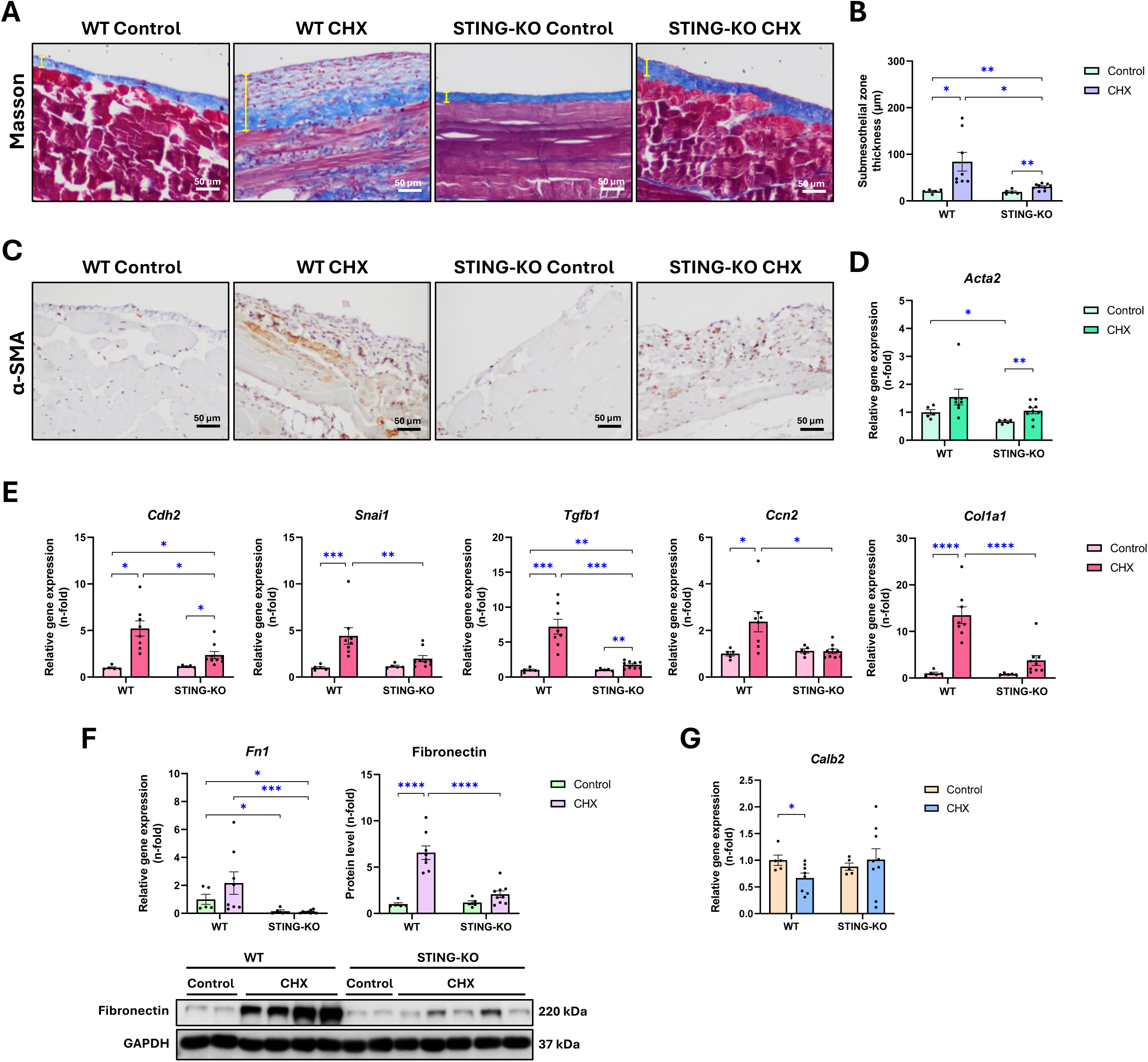
Peritoneal membrane thickening and fibrosis levels in WT and STING-KO mice exposed to CHX for 30 days. Male C57BL/6J wild-type (WT) and STING-deficient (STING-KO) mice were intraperitoneally injected with 0.1% chlorhexidine gluconate (CHX) for 30 days. **(A-C)** Parietal peritoneal tissue sections were used for Masson’s trichrome staining **(A,B)** and α-SMA immunostaining **(C)**. Microscopy images correspond to a representative animal from each group. The yellow lines in Masson images indicate the width measured. Scale bar: 50 μm. **(D-G)** Relative gene expression of α-SMA (*Acta*) **(D),** other mesothelial-to-mesenchymal transition and fibrosis markers **(E)**, fibronectin (*Fn1*) **(F)**, and the mesothelial marker calretinin (*Calb2*) **(G).** Relative gene expression was analyzed by RT-qPCR from total RNA of parietal peritoneal tissue, using Gapdh as a housekeeping gene. **(F)** Fibronectin protein levels assessed by western blotting from total parietal peritoneum proteins, using GAPDH as loading control. Results are expressed as fold change (n-fold) compared to the control group and are represented as the mean ± SEM of 5–9 animals per group. * *p* < 0.05 and ** *p* < 0.01; *** *p* < 0.001; and **** *p* < 0.0001.

Peritoneal transcriptional levels of proinflammatory genes in WT mice, including cytokines, chemokines, macrophage markers, IFNs, and ISGs, remained upregulated after 30 days of CHX exposure (Figure 6A). However, in the STING-deficient mice exposed to CHX, no alterations were observed in most components of this gene signature when compared to control mice (Figure 6A). Peritoneal NF-κB pathway activation was also prevented by STING deficiency, as indicated by the prevention of NF-κB inhibitor α (IκBα) (Figure 6B) and p65/NF-κB (Figure 6B and 6C) phosphorylation. In addition, the recruitment of cells into the peritoneal cavity in response to CHX was quantified by flow cytometry. There was a notable reduction in the total number of recruited cells in STING-KO mice relative to WT mice (Figure 6D). CD3^+^ T lymphocytes (CD4^+^ T helper and CD8^+^ T cytotoxic), neutrophils (CD11b^+^ F4/80^-^ Ly6G^+^), and macrophages (CD11b^+^ F4/80^+^), both peritoneal tissue-resident (CD11b^+^ F4/80^high^) and blood monocyte-derived (CD11b^+^ F4/80^low^) were among the recruited cell subsets displaying a lowered peritoneal transit in STING-KO mice. In line with this restrained recruitment of immune cells towards the peritoneal cavity, inflammatory cell infiltration into the peritoneal tissue was also reduced in CHX STING-KO mice (Supplementary Figure S4). Together with the increased peritoneal inflammation induced by CHX, we found a rise of the oxidative stress and the antioxidant response in WT mice, including higher gene expression levels of the pro-oxidant enzymes *Nox1* and *Nox4* (Supplementary Figure S5A), increased transcription and activation of the nuclear factor erythroid 2-related factor 2 (NFR2; detected through *Nfe2l2* gene expression and p-NRF2 nuclear localization, respectively) (Supplementary Figure S5B and 5D,), and overexpression of NRF2 antioxidative target enzymes, such as catalase (*Cat*), heme oxygenase 1 (*Hmox1*), and superoxide dismutases (*Sod1* and *Sod2*) (Supplementary Figure S5). All these markers of the oxidative stress response were decreased in the peritoneum of CHX-exposed STING-KO mice versus CHX-exposed WT mice. Altogether, these findings demonstrate that STING absence ameliorates peritoneal fibrosis, inflammation, and oxidation induced by chronic and repeated exposure to CHX in a model of advanced peritoneal damage.

**Figure 6.**
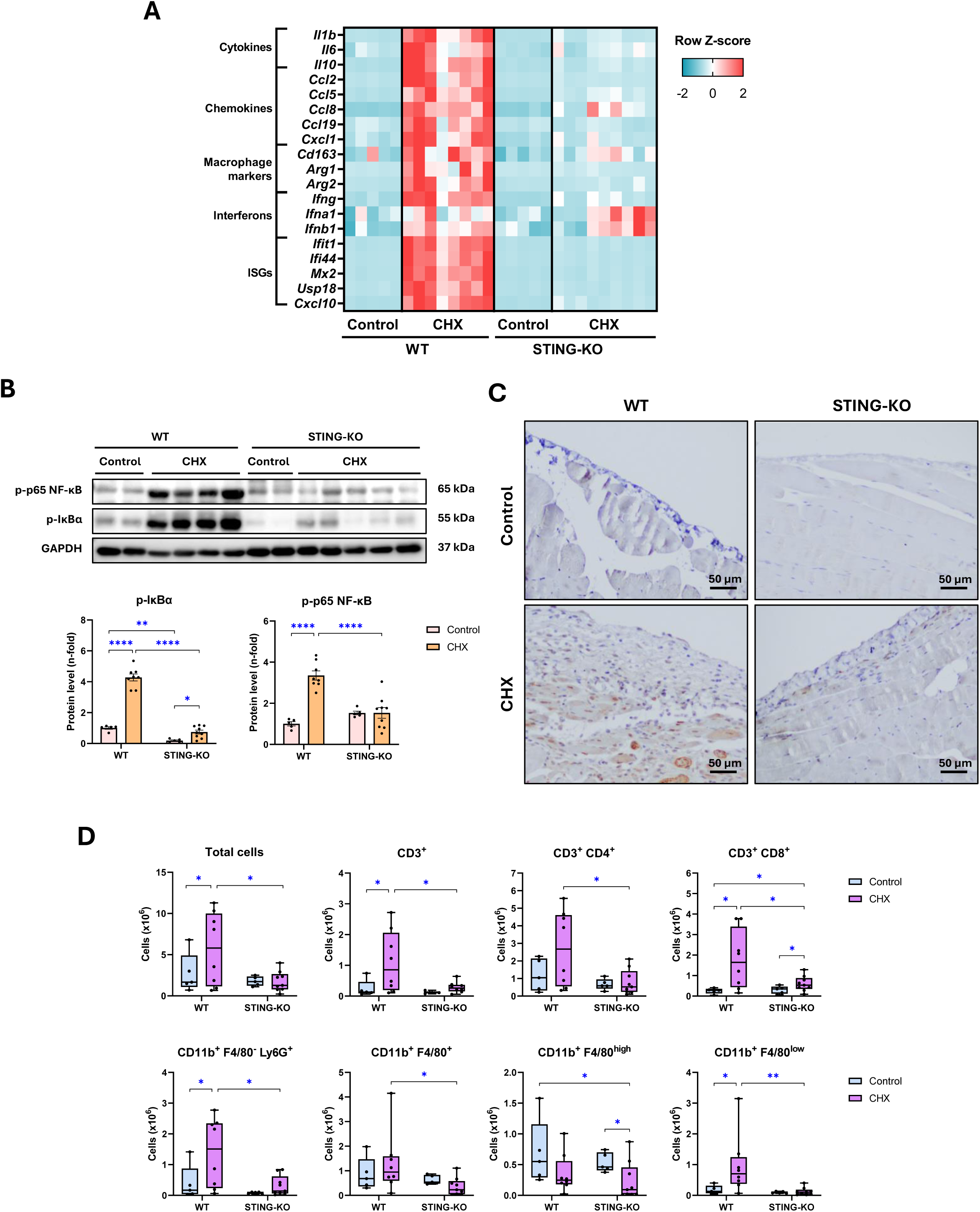
Peritoneal inflammation evaluation in WT and STING-KO mice exposed to CHX for 30 days. Male C57BL/6J wild-type (WT) and STING-deficient (STING-KO) mice were intraperitoneally injected with 0.1% chlorhexidine gluconate (CHX) for 30 days. **(A)** Heatmap represents the row *Z*-score of the relative expression levels of inflammatory markers. Relative gene expression was analyzed by RT-qPCR from total RNA of parietal peritoneal tissue, using *Gapdh* as housekeeping gene. **(B)** Protein levels of phosphorylated p65 subunit of NF-κB (p-p65 NF-κB) and phosphorylated IκBα (p-IκBα) assessed by western blot from total proteins of parietal peritoneal tissue, using GAPDH as loading control. **(C)** Immunohistochemical detection of the p-p65 NF-κB protein (brown) in parietal peritoneal tissue sections. Microscopy images correspond to a representative animal from each group. Scale bar: 50 μm. **(D)** Flow cytometry in peritoneal lavages from mice to identify cells present in the peritoneal cavity. Peritoneal cells were labeled with specific antibodies tagged with different fluorophores to identify CD3^+^, CD4^+^, CD8^+^, CD11b^+^, Ly6G^+^, and F4/80^+^ cells. Western blot results are expressed as fold change (n-fold) relative to the control group and are represented as mean ± SEM. Cytometry results are represented in scattered box plots with min-to-max whiskers, and the median and quartiles are shown. N = 5-9 per group. * *p* < 0.05; ** *p* < 0.01; and **** *p* < 0.0001.

### *In vitro* pharmacological targeting of STING inhibits proinflammatory responses in MCs and macrophage-mediated MMT

MCs can undergo MMT, acquiring myofibroblast phenotype and contributing to ECM accumulation and fibrosis [33] and to the inflammatory response by secreting a variety of cytokines [7,34]. To evaluate the impact of pharmacological STING inhibition, MeT-5A MCs were pretreated with the covalent STING inhibitor C-176 (1 μM) and then stimulated with the master profibrotic factor TGF-β (3 ng/mL) for 48 hours. No significant changes were observed in the TGF-β-induced gene expression levels of MMT and fibrosis markers after C-176 treatment (Figure 7A). Despite this, the STING pathway remained sensitive in Met-5A cells when challenged with 5 ng/mL TNF-α for 48 h, as judged by the induced gene expression of chemokines (*CCL2* and *CCL5*) and ISGs (*CXCL10* and *USP18),* which was however dampened in the presence of C-176 (Figure 7B). Thus, these results suggest that STING does not directly induce MMT in MCs but contributes to the maintenance of the inflammatory response. Since STING-positive macrophages were noticed *in vivo* (Figure 4B), we wanted to know whether STING inhibition on macrophages influences MMT in MCs. To this aim, peritoneal macrophages from mice were treated with LPS in the presence or the absence of C-176 to obtain macrophage-derived conditioned medium (MCM). After that, MeT-5A MCs were growth in presence of MCM from LPS-activated macrophages (with or without C-176 pre-treatment) (Figure 7C). In the LPS-activated peritoneal macrophages, C-176 significantly decreased inflammatory gene expression (Figure 7D). In MCs, LPS-MCM induced a significant increase in the gene expression of MMT and profibrotic markers, suggesting that macrophages secrete factors that modulate MCs. By contrast, the profibrotic effect of the LPS-MCM from C-176 pre-treated macrophages was significantly lower (Figure 7E). These results indicate that *in vivo* peritoneal MMT and fibrosis may be, at least in part, mediated by the activation of STING in macrophages.

**Figure 7.**
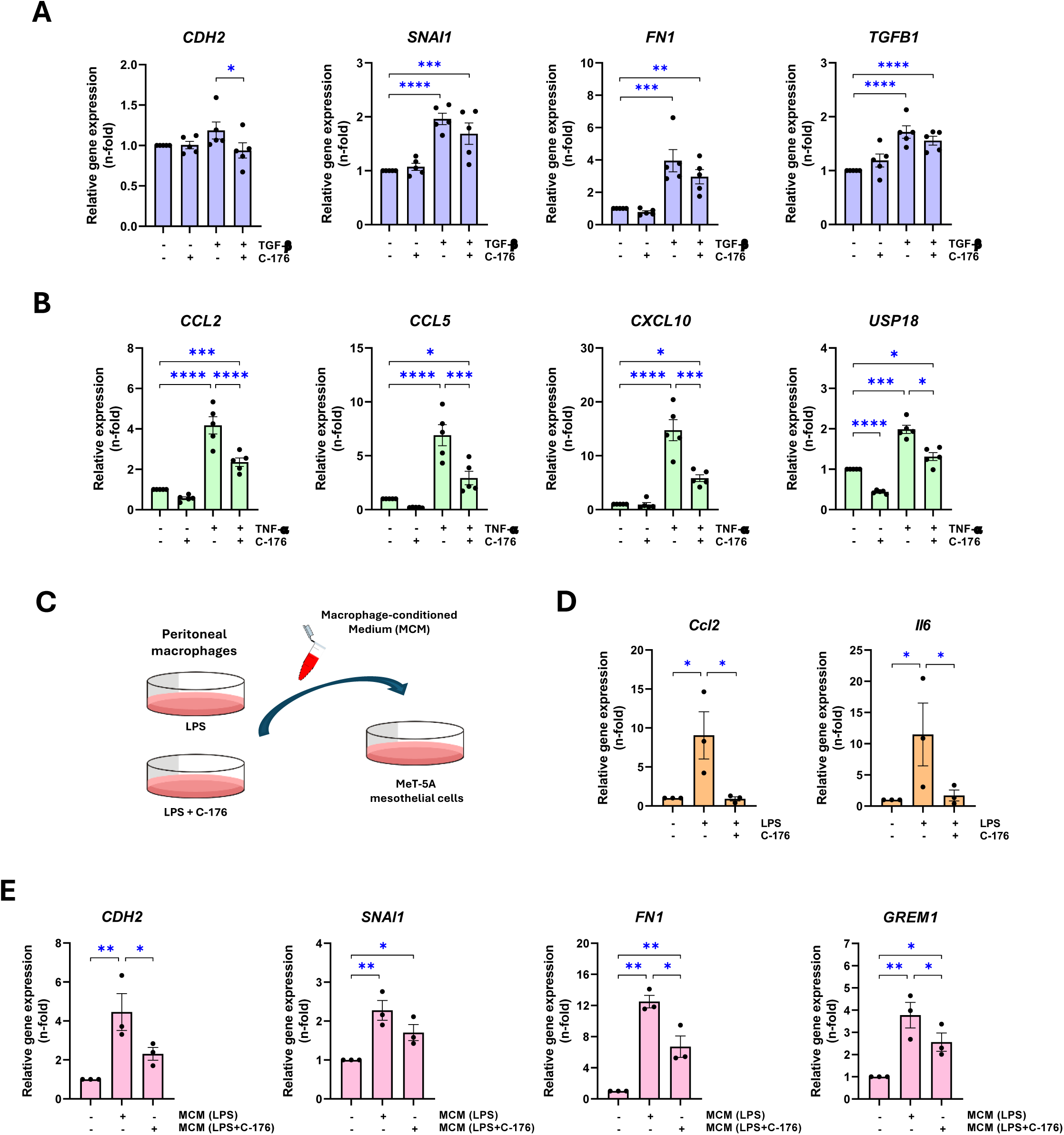
Pharmacological inhibition of STING in cultured mesothelial cells and macrophages. **(A, B)** Relative gene expression of mesothelial-to-mesenchymal transition/fibrosis (*CDH2, SNAI1, FN1,* and *TGFB1*) and inflammatory (*CCL2* and *CCL5*) and interferon-stimulated (*CXCL10* and *USP18*) genes in human MeT-5A mesothelial cells pretreated with 1 µM C-176 for 1 hour and then stimulated with recombinant TGF-β (3 ng/mL) **(A)** or TNF-α (5 ng/mL) **(B)** for 48 hours. **(C)** Schedule of the in vitro study design for the assessment of macrophage-mesothelial cell interaction. Mouse peritoneal macrophages were pretreated with 1 µM C-176 for 1 h and then activated with LPS (1 µg/mL). After 6 h of incubation, the medium was changed and incubated for 24 h to obtain macrophage-conditioned medium (MCM) which was used as a stimulus to MET-5A cells. **(D)** Relative gene expression of inflammatory markers in cultured macrophages challenged with LPS in the presence of the absence of C-176. **(E)** Relative gene expression of mesothelial-to-mesenchymal transition and fibrosis markers in MeT-5A cells incubated for 48 h with MCM from LPS-activated macrophages with or without C-176 treatment. Relative gene expression levels of were analyzed by RT-qPCR from total RNA, using human *GAPDH* or mouse *Gapdh* as housekeeping gene, as appropriate. Results are expressed as fold change (n-fold) relative to the control condition (first column) and represented as the mean ± SEM of 3-5 independent experiments. * p < 0.05; ** p < 0.01; *** p < 0.001; and **** p < 0.0001.

### *In vivo* pharmacological inhibition of STING ameliorated CHX-induced peritoneal damage

To test the *in vivo* potential protective effect of STING pharmacological inhibition in peritoneal damage, we developed a 10-day CHX exposure model in WT mice together with daily intraperitoneal administration of C-176 (750 nmol/mice, dissolved in corn oil as vehicle). Vehicle administration significantly enhanced CHX-induced peritoneal damage (Figure 8), when compared with CHX-only exposure. Despite the harmful effect of vehicle, C-176 treatment was able to significantly attenuate CHX-induced PM thickening (Figure 8A) and *Tgfb1* and *Col1a2* gene expression levels (Figure 8B), in comparison to CHX + vehicle group. Moreover, C-176 treatment significantly reduced macrophage cell infiltration into the peritoneal tissue induced by CHX + vehicle peritoneal exposure (Figure 8C), together with normalization in the expression levels of a panel of inflammatory genes, including chemokines, cytokines, macrophage markers and ISGs (Figure 8D). Taken together, our results demonstrate that pharmacological inhibition of STING improves the peritoneal injury caused by CHX in mice.

**Figure 8.**
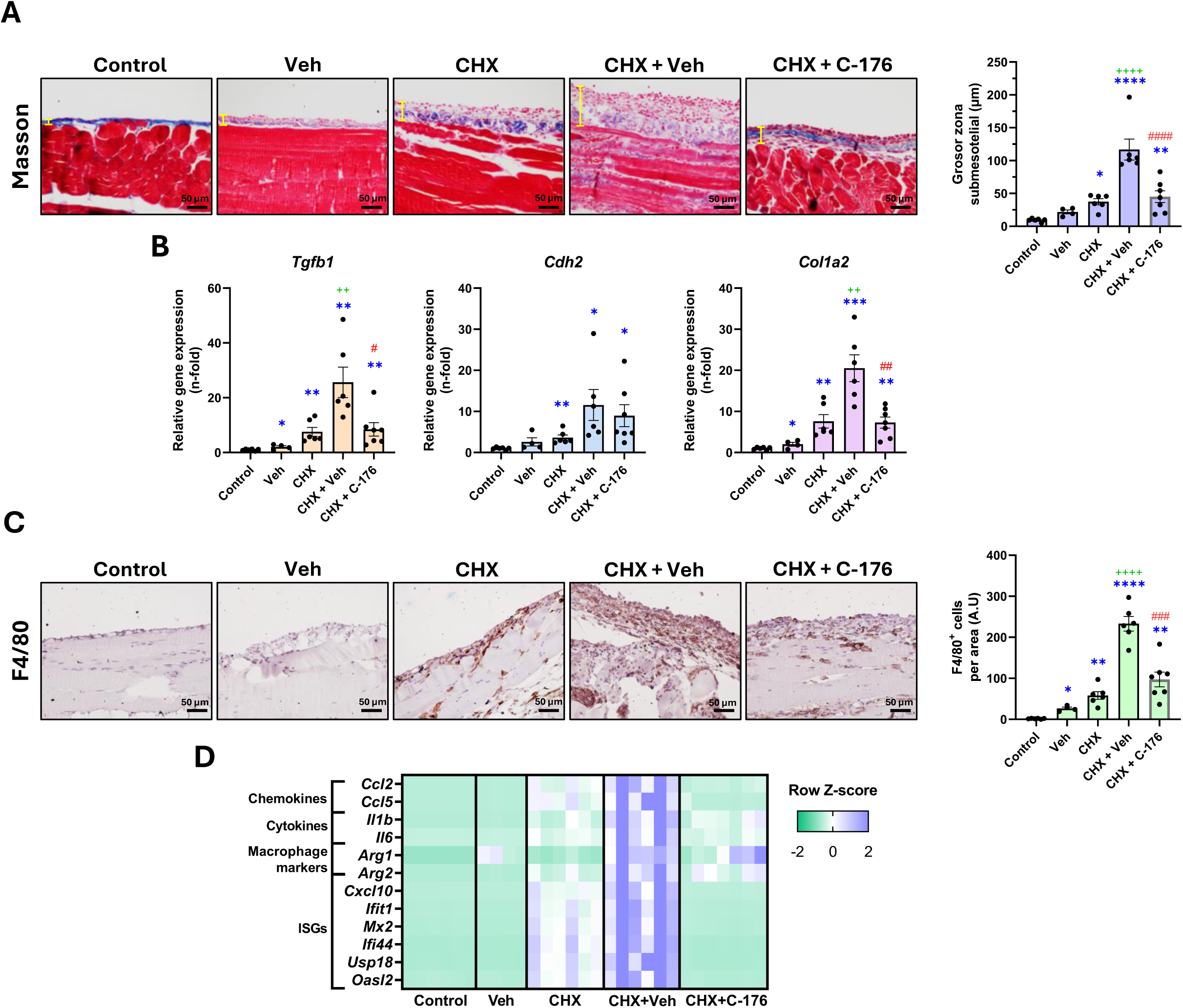
Pharmacological inhibition of STING in CHX-infused mice. Male C57BL/6J wild-type mice were intraperitoneally injected with 0.1% chlorhexidine gluconate (CHX) together with pharmacological STING inhibitor C-176 or its vehicle (Veh, corn oil) for 10 days. Untreated and vehicle-treated mice were used as controls. **(A)** Peritoneal membrane thickness assessment. The figure shows Masson’s trichrome stained parietal peritoneal tissue sections of representative mice from each group (left) and the corresponding quantification of submesothelial zone thickness (right). The yellow lines indicate the thickness measured. Scale bar: 50 μm. **(B)** Relative gene expression of markers of mesothelial-to-mesenchymal transition and fibrosis markers. **(C)** Immunohistochemical staining of F4/80 to identify infiltrating macrophages in peritoneal tissue. Microscopy images correspond to a representative animal from each group (left) and the corresponding quantification of F4/80^+^ cells (right) are shown. Scale bar: 50 μm. **(D)** Heatmap represents the row *Z*-scores of the level of expression of inflammatory markers. Relative gene expression was analyzed by RT-qPCR from total RNA of parietal peritoneal tissue, using *Gapdh* as housekeeping gene, and expressed as fold change (n-fold) relative to control. Results are represented as mean ± SEM of 4–7 animals per group. * *p* < 0.05; ** *p* < 0.01; *** *p* < 0.001; and **** *p* < 0.0001 compared to Control group. ++ *p* < 0.01 and ++++ *p* < 0.0001 compared to CHX group. # *p* < 0.05; ### *p* < 0.001; and #### *p* < 0.0001 compared to CHX + Veh group. A.U.: arbitrary units.

### STING deficiency alleviates post-surgical peritoneal adhesions

Intra-abdominal adhesions also occur in PD patients [35,36]. To study the role of STING in the development of adhesions, we performed a peritoneal adhesion model of ischemic buttons (IBs) surgically made in the abdominal wall of WT and STING-KO mice. In WT mice, we found abundant STING-positive cells in peritoneal areas close to IBs and in the peritoneum-organ adhesion interface (Supplementary Figure S6A and S6B). Moreover, the protein levels of STING, TBK1, and IRF3 were found increased in IB tissue compared to control peritoneal tissue (Supplementary Figure S6B). In STING-deficient mice, the extent of the IB covered by the adhesion (grade) tends to be less than that observed in WT mice and the tenancy and total score of the adhesions were significantly decreased (Figure 9A). Infiltration of macrophages (Figure 9B) and T lymphocytes (Supplementary Figure S7) was also reduced in IBs from STING-deficient mice compared to WT mice. In addition, the gene expression levels of the inflammatory and ISG chemokine *Cxcl10* and other ISGs (*Mx2, Ifit1,* and *Usp18*) were reduced in the IBs from STING-KO mice in comparison with WT mice. Altogether, these results show that STING deficiency ameliorates peritoneal adhesions in mice.

**Figure 9.**
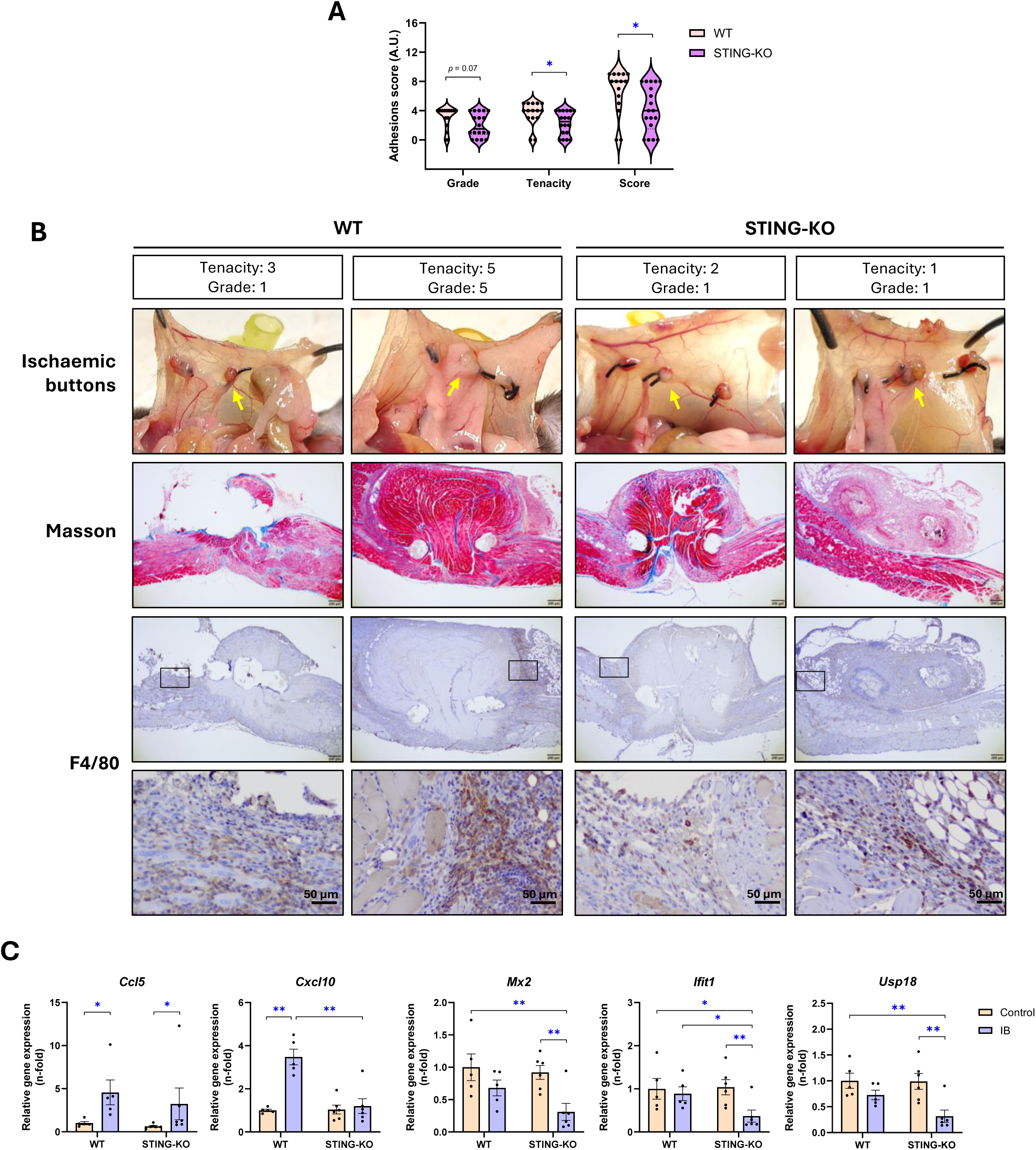
Comparative evaluation of the grade, tenacity, and inflammation of peritoneal adhesions in WT and STING-KO mice. Three ischemic buttons (IBs) were surgically made in the peritoneum of male C57BL/6J wild-type (WT) and STING-deficient (STING-KO) mice. Adhesion formation on the IBs was assessed 5 days after surgery. **(A)** Peritoneal adhesion grade, tenacity and total score values are represented by violin scatterplots showing the median, quartiles, distribution and density of the data for each experimental group (n = 5-6 mice per group, 3 IBs/mouse). **(B)** Macroscopic and microscopic analyses of adhesions formed on IBs surgically made in mice. Yellow arrows in top panel indicate the IB to which correspond the lower Masson’s trichrome (middle panel) and F4/80 immunostaining (bottom panels) micrographs. Squared areas encompassing adhesion interfaces and F4/80^+^ cells are shown at a higher magnification below. Microscopy images correspond to two representative animals from each group. Scale bars: 200 and 50 μm. **(C)** Relative expression of inflammatory and interferon-stimulated genes analyzed by RT-qPCR from total RNA of parietal peritoneal tissue, using *Ppia* as housekeeping gene. Results are expressed as fold change (n-fold) relative to the control group and represented as the mean ± SEM of 5-6 animals per group. * *p* < 0.05 and ** *p* < 0.01.

### STING deficiency alleviated *S. epidermidis*-induced peritonitis

Peritonitis is an infection within the abdominal cavity primarily caused by bacteria that often impairs PD [37]. *Staphylococcus epidermidis* is one of the most frequently isolated causes of PD-associated peritonitis [38,39]. To evaluate the participation of STING in peritonitis, we induced peritonitis in WT and STING-KO mice by a single intraperitoneal injection of live *S. epidermidis* (5 x 10^8^ cfu/mice) followed by the peritoneal tissue examination after 72 hours. *S. epidermidis* peritoneal exposure induced a significant increase in PM thickness in both WT and STING-KO mice, but no significant differences were observed between these two groups (although a tendency to decrease can be observed in STING-KO mice) (Figure 10A). Regarding inflammation, STING deficiency was able to prevent *S. epidermidis*-induced upregulation of inflammatory genes such as chemokines and cytokines (including the ISG *Cxcl10*, Figure 10B), macrophage markers *Arg1* and *Arg2* (Figure 10C), and type I interferons *Ifna1* and *Ifnb1* (Figure 10D). In addition, STING-KO mice showed decreased *S. epidermidis*-induced IκB phosphorylation, suggesting less NF-κB pathway activation in comparison with WT mice (Figure 10E). Thus, STING deficiency improved peritoneal inflammation induced by *S. epidermidis* in mice.

**Figure 10.**
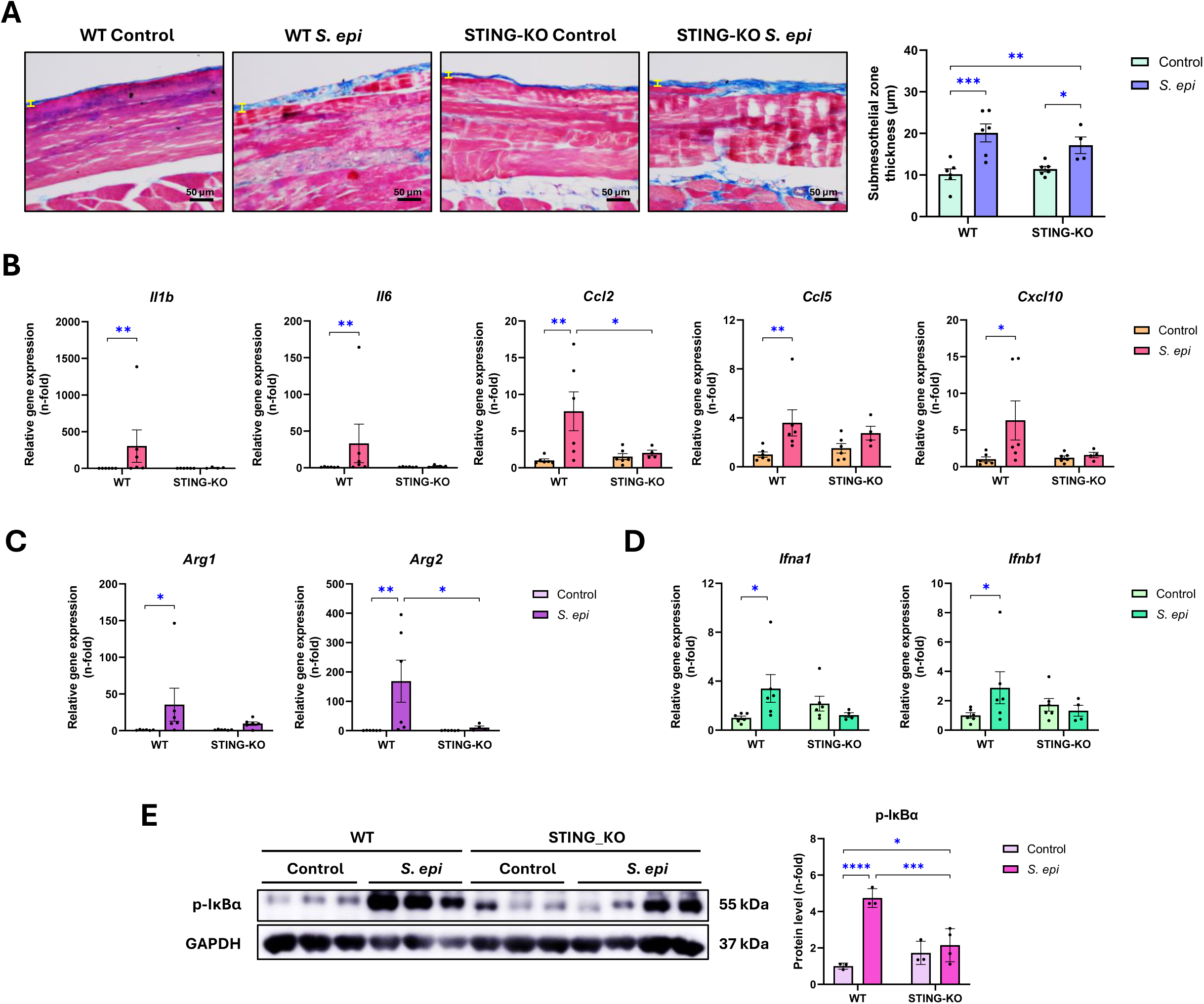
Inflammatory response in *S. epidermidis*-induced peritonitis in WT and STING-KO mice. Female C57BL/6J wild-type (WT) and STING-deficient (STING-KO) mice were injected with a single dose of live *S. epidermidis* bacteria (*S. epi*, 5 × 10^8^ cfu/mouse) and evaluated 72 hours after injection. **(A)** Peritoneal membrane thickness assessment. The figure shows Masson’s trichrome stained parietal peritoneal tissue sections of representative mice from each group (left) and the corresponding quantification of submesothelial zone thickness (right). The yellow lines indicate the width measured. Scale bar: 50 μm. **(B-E)** Relative gene expression of cytokines and chemokines **(B)**, macrophage markers **(C)**, and type I interferons **(D)** analyzed by RT-qPCR from total RNA of parietal peritoneal tissue, using *Ppia* as housekeeping gene. **(E)** Protein levels of phosphorylated IκBα (p-IκBα) assessed by western blot from total proteins of parietal peritoneum, using GAPDH as loading control. Gene and protein level results are expressed as fold change (n-fold) relative to the control group. All results are represented as the mean ± SEM of 3–6 animals per group. * *p* < 0.05; ** *p* < 0.01; *** *p* < 0.001; and **** *p* < 0.0001.

## Discussion

In this article, we describe for the first time that STING is expressed in the injured human peritoneum and that STING activation plays a role in several preclinical models of PD-associated damage, including peritoneal exposure to chemical agents (CHX and PDF), peritoneal adhesions, and bacterial peritonitis. Genetic deletion of STING or its pharmacological blockade diminished peritoneal inflammation and fibrosis, showing the contribution of this pathway to the initiation and progression of peritoneal damage in response to sterile and nonsterile aggressions that engage innate immune responses against the cytoplasmic DNA accumulation. In light of these findings, we propose that STING represents a promising novel therapeutic target for preventing PD-associated peritoneal deterioration.

PD is a life-saving therapy for ESKD patients, however, repeated and prolonged exposure to PDFs induces peritoneal damage and other related complications, such as bacterial infections or catheter adhesions that worsen ultrafiltration capabilities of the peritoneal membrane and lead to therapy discontinuation [40]. Previous transcriptomic studies using peritoneal biopsies from PD and uremic patients without a history of PD have shown functional enrichment in pathways and gene networks associated with inflammation and the immunological response [41]. Moreover, RNA-seq studies in children with established PD have confirmed the involvement of inflammatory, immunologic, and stress responses [42]. Consistent with the notion that inflammation and the immune response are key drivers of PD injury, our study in a preclinical model resembling chronic PD exposure induced by CHX has revealed that the most upregulated genes in response to CHX-induced peritoneal damage are targets of inflammatory pathways, including chemokines (*Ccl2* and *Ccl7*), cytokines (*Il1b*), and ISGs (*Cxcl10* and *Isg15*). In addition, functional analysis from the peritoneal transcriptomic data showed that among the most enriched terms were innate immune-related pathways, including one not previously described, the cytosolic DNA-sensing pathway, and others already known in PD-associated damage, such as those relaying on NLRs, TLRs, NF-κB and TNF-α.

We and other researchers have recently focused on identifying new mediators of the immune response involved in PD injury and their potential targeting to reduce peritoneal deterioration [7]. Accordingly, preclinical studies have shown that blockade of several proinflammatory pathways ameliorated peritoneal damage, as shown by inhibition of TLRs [43], TNF family proteins [44], IL-17A [45], JAK2/STAT3 [46], CCL8/CCR8 [47], and NF-κB [8], among others. In contrast, no data about modulating STING has been described to date.

DAMPs are key pro-inflammatory molecules that are normally prevented from entering the extravesicular or extracellular space, but when exposed they initiate the innate immune response. Among DAMPS, mitochondrial and genomic double-stranded DNA (dsDNA) accumulated in the cytoplasm of stressed/damaged cells has recently emerged as a main driver of autoinflammatory, autoimmune and acute or chronic illnesses, including cancer [48–50]. This cytosolic DNA is recognized through several sensors and adapter proteins belonging to the intracellular DNA sensing pathway. STING has arisen as a conspicuous intracellular component of the DNA-sensing pathway whose excessive, unrestrained, or extemporaneous activation, may result in pathological inflammation and tissue damage [51–53]. Together with the IFN-I pathway, which we found to be activated by STING-dependent TBK1/IRF3 phosphorylation and further ISG induction, NF-κB-mediated inflammation is evolutionarily conserved and has been implicated in STING-mediated responses, including antiviral and antitumor activities [54,55]. STING recruits both TBK1 and IKKε to activate the NF-κB pathway and induce the transcription of a series of pro-inflammatory cytokines and chemokines in immune and epithelial cells [17,20,56]. In this regard, the bioinformatic analysis of the RNA-seq dataset from our peritoneal damage model predicted proinflammatory gene programs under the transcriptional control of NF-κB. Previous studies have shown that inhibition of NF-κB with parthenolide, a widely used IKKα/β inhibitor compound, ameliorated peritoneal damage by inhibiting inflammation and fibrosis [8,57]. Remarkably, we showed that in murine peritoneal tissue damaged by CHX, NF-κB was blunted in STING-deficient mice as well as by the pharmacological blockade of the palmitoylation-induced STING activation [58]. Indeed, treatment with C-176 restrained the synthesis of NF-κB-dependent cytokines and hence immune cell infiltration in the CHX-exposed peritoneum. Studies in other pathologies confirm our data, as observed with the pharmacological STING inhibition by H-151, which alleviated NF-κB-dependent inflammatory responses in a model of dermatitis, reducing the secretion of inflammatory factors such as IL-6 and TNF-α by keratinocytes and macrophages [59]. Moreover, genetic deletion of STING significantly ameliorates hyperglycemia-induced oxidative and mitochondrial endothelial damage, resulting in cGAS-STING-dependent NF-kB activation and inflammation in diabetic mice [60]. These antecedents and our results indicate that STING plays a pivotal role in promoting the progression of NF-κB-induced peritoneal injury, suggesting that it may be a promising novel therapeutic target to prevent peritoneal deterioration. On the other hand, the peritoneal damage induced by infectious agents is mainly mediated by pathogen-associated molecular patterns (PAMPs) and their immune receptors, which include TLRs and NLRs, that coordinate the elimination of pathogens and infected cells by the production of inflammatory mediators [61]. TLRs expressed by peritoneal macrophages and mesothelial cells contribute to peritoneal infection [25]. Our RNA-seq dataset, although done in sterile inflammation, also showed the activation of TLRs and NLRs pathways together with the STING pathway. Moreover, in our model of peritonitis induced by live *S. epidermidis* administration, STING-deficient mice showed decreased NF-κB activation leading to downregulation of dependent chemokines (*Ccl2* and *Ccl5*), cytokines (*Il1b* and *Il6*) and macrophage markers (*Arg1* and *Arg2*), indicating that inflammation promoted from the STING/NF-κB pathway also participates in peritoneal infection.

The systemic pathological consequences of PD-associated peritonitis and its control are currently challenging issues. In ESKD patients under PD, bacterial peritonitis is common and has been associated with an increased risk of cardiovascular mortality [62,63]. Therefore, anti-inflammatory therapies administered during or after PD-associated peritonitis combined with standard anti-microbial therapies have been proposed to reduce long-term cardiovascular risk [64]. Several studies have shown the involvement of the STING pathway inactivation in ameliorating cardiovascular diseases, either through genetically deficient mice for cGAS, STING, IRF3, or the type I IFN receptor IFNAR, and by direct pharmacological blockade of STING (C-176 or H-151) or the STING-dependent IRF3 recruitment in the signalosome (Astin C) [65–71]. On the other hand, in a murine model of bacterial peritonitis, treatment with an inhibitor of Calprotectin, a DAMP and TLR4 ligand, robustly inhibited the short and long-term systemic and vascular inflammatory consequences of peritonitis, without affecting the bacterial clearance [27]. The CANTOS trial and others using monoclonal antibodies to IL-1β demonstrated that the reduction of chronic inflammation effectively reduces cardiovascular risk, particularly in CKD patients. However, this strategy was associated with elevated risk of severe infections, which can ultimately result in mortality [72,73]. This body of evidence suggests that future studies investigating combined anti-inflammatory treatments, such as TLR4 or STING inhibitors, especially in populations prone to infections, are a likely pharmacological alternative to address the cardiac consequences of chronic PD and peritonitis associated with renal failure.

Peritoneal fibrosis is an undesirable consequence of chronic exposure to PDFs, characterized by an excessive accumulation of ECM proteins, leading to progressive thickening of the submesothelial compact zone and membrane dysfunction [74]. Biochemical mediators and biomechanical forces converge in the activation of TGF-β1 to trigger signals leading to MMT [75–77], mainly by the Smad pathway activation [78–80], although other signaling mechanisms has also been escribed, such as MEK-ERK1/2-Snail-1 [81] and CAV1-YAP/TAZ pathway [82]. STING has been described as an essential regulator of fibrosis in human and mouse hypertrophic hearts [83] and experimental kidney, lung, and liver fibrosis [84]. Studies using conditional *Pdk1*-and *Cgas*-knockout mouse models demonstrated that STING decreases cystogenesis and renal fibrosis [85]. In contrast, STING deficiency was also found to lead to increased lung fibrosis in an unexpected type I IFN-independent manner [86]. The protective role of STING against fibrosis has also been shown in a model of chronic pancreatitis [87]. A key feature of our study shows that targeting STING (by either genetic or pharmacological inhibition) decreases peritoneal fibrosis in CHX-induced damage by regulating the expression of fibrotic inducers, such as TGF-β1 and restoring the expression of mesothelial markers, therefore preventing MMT, a key process in peritoneal fibrosis. Overall, these data suggest that STING, despite modulating the fibrotic response in a disease-specific manner, promotes peritoneal fibrosis, and therefore, its targeting may help to improve the pathological fate during PD.

We have previously characterized the transcriptomic reprogramming of the mesothelial cells to myofibroblast-like phenotype defining a panel of pro-MMT markers [88]. The *in vitro* studies done here demonstrates that STING does not act as a direct inducer of MMT in MCs. However, STING plays a role in the inflammatory response in this cell type, as evidenced by the inhibition of TNFα-induced proinflammatory gene upregulation by C-176. Thus, these data confirm the key role of the STING signaling in sustaining chronic inflammation and that this proinflammatory loop contributes to peritoneal damage progression.

STING is expressed in a wide variety of tissues and cells, including both immune and resident cells [53]. In human peritoneal biopsies positive STING cells were found in the submesothelial compact zone in fibrotic areas. Moreover, in CHX-treated mice, positive infiltrating STING cells, mainly macrophages, were found. In our RNA-seq experiment, bioinformatic tools and databases reveal that macrophages were the most enriched cell subset in CHX-induced peritoneal damage. Accordingly, the presence of infiltrating macrophages was found in the submesothelial compact zone of the CHX-damaged peritoneum, associated with the upregulation of proinflammatory genes, including chemokines involved in macrophage recruitment as CCL2. Analysis of peritoneal effluent from PD patients has shown a correlation between the degree of fibrosis and the presence of CCL18/CD163 expressing macrophages, which phenotypically and functionally resemble polarized M2 macrophages *in vitro* [89]. However, the role of macrophages during PD-induced peritoneal damage depends on the pathological context and macrophage subtype [7]. Recently, macrophage reprogramming and immunometabolism have been linked to the activation of STING [90]. In CHX-induced peritoneal damage, the number of infiltrating macrophages was significantly diminished by STING deficiency or pharmacological inhibition. Moreover, in the other models of peritoneal damage studied here, bacterial peritonitis and peritoneal adhesions, the inhibition of STING blunted macrophage infiltration and proinflammatory genes upregulation. Furthermore, the identification of macrophages in the submesothelial compact zone, together with peritoneal thickness and fibrosis in CHX-treated mice, suggest a link between STING activity in macrophages and fibrosis/MMT. Our *in vitro* experiments on MCs cultured in the presence of conditioned medium from LPS-activated macrophages treated with a pharmacological STING inhibitor demonstrate that activation of the STING pathway in macrophages is involved in the induction of MMT. These data show a crosstalk between the immune system and cells of the peritoneal stroma in the fibrotic process and suggest that *in vivo* peritoneal MMT and fibrosis may be, at least in part, mediated by the activation of the STING pathway in macrophages.

Surgical procedures also cause significant damage to the peritoneum. In some cases, this leads to the development of peritoneal adhesions, which are composed of fibrous bands that connect organs to each other or to the parietal peritoneal wall. Peritoneal adhesions represent a major cause of postoperative morbidity and pose a significant challenge to public health [91]. Patients with intra-abdominal adhesions undergoing PD are at an increased risk of developing abdominal pain or catheter occlusion [36]. A rare complication of long-term PD is encapsulating peritoneal sclerosis, which is characterized by fibrosis and adhesions of the peritoneum to the small bowel loops, resulting in intestinal obstruction and fatal consequences [92]. Accordingly, STING-deficient mice have fewer peritoneal adhesions, demonstrating that STING inhibition can not only prevent early inflammation but also its chronic consequences.

In conclusion, our study identifies STING as a key regulator of peritoneal injury progression through induction of inflammation, macrophage recruitment and fibrosis. Our results highlight a potential therapeutic strategy of inhibiting STING activation with pharmacological agents, such as C-176, for the treatment of peritoneal injury.

## Acknowledgements

We thank Irene Rubio for her technical support, especially in the processing of histological samples.

## Author contributions

V.M., G.T.G.-M., A.-C.R., M.L.-C., A.M.R., and M.R.-O. conceived and designed the work. V.M., J.G.-G., G.T.G.-M., R.R., P.S., L.T.-S., and S.R.-M. performed the experiments and obtained the data. V.M., G.T.G.-M., P.S., R.R., J.J.-H., A.M.R., and M.R.-O. analyzed and interpreted the results. V.M., A.M.R., and M.R.-O. drafted the manuscript. V.M., A.M.R., A.C.-R., M.L.-C., A.O., and M.R.-O. edited, and revised the manuscript. All authors approved the final version of the manuscript.

