## Supplementary Figure for "Targeting STING in experimental peritoneal damage: a novel approach in peritoneal dialysis therapy"

### Suppl. Figure S1

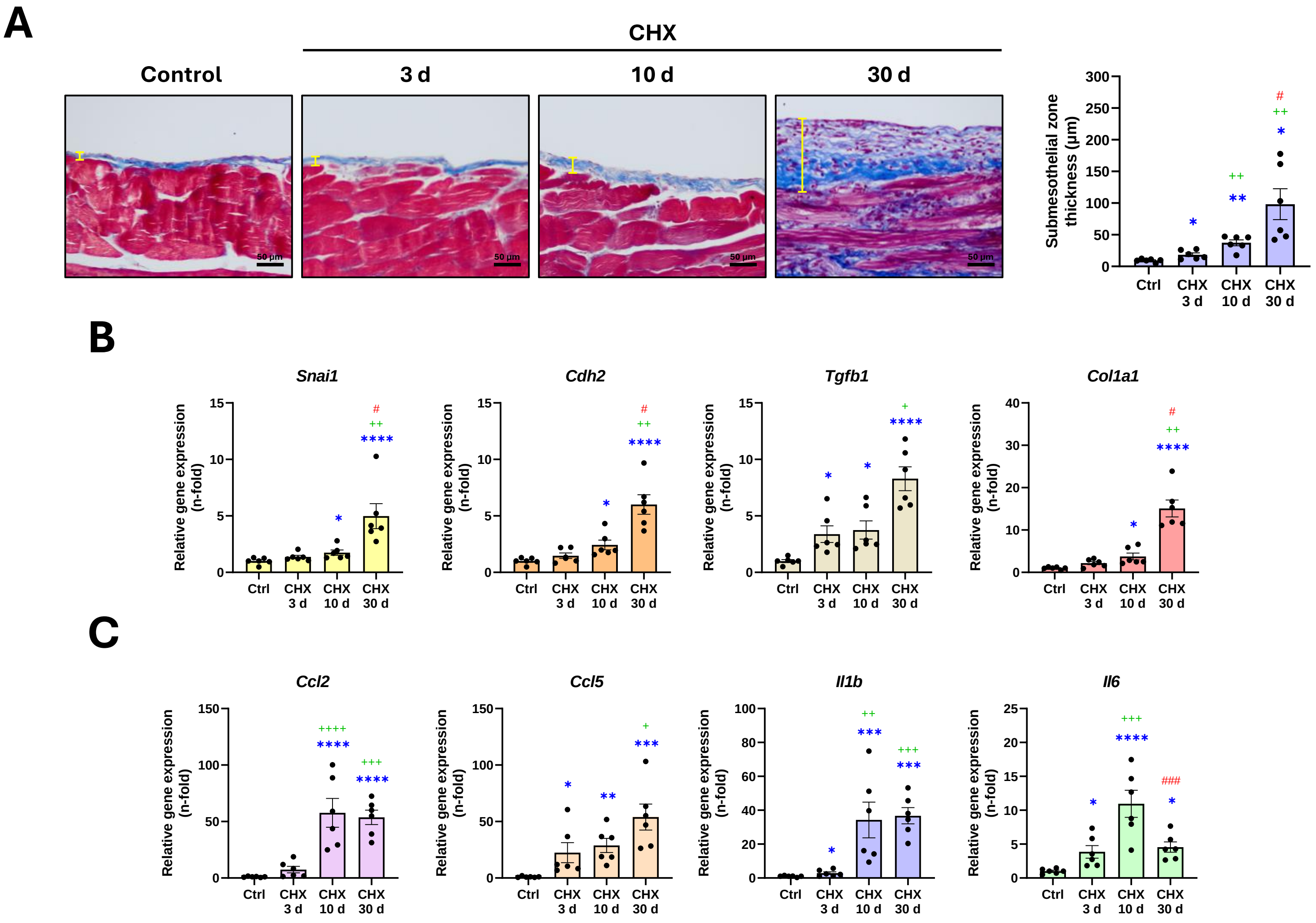

### Suppl. Figure S2

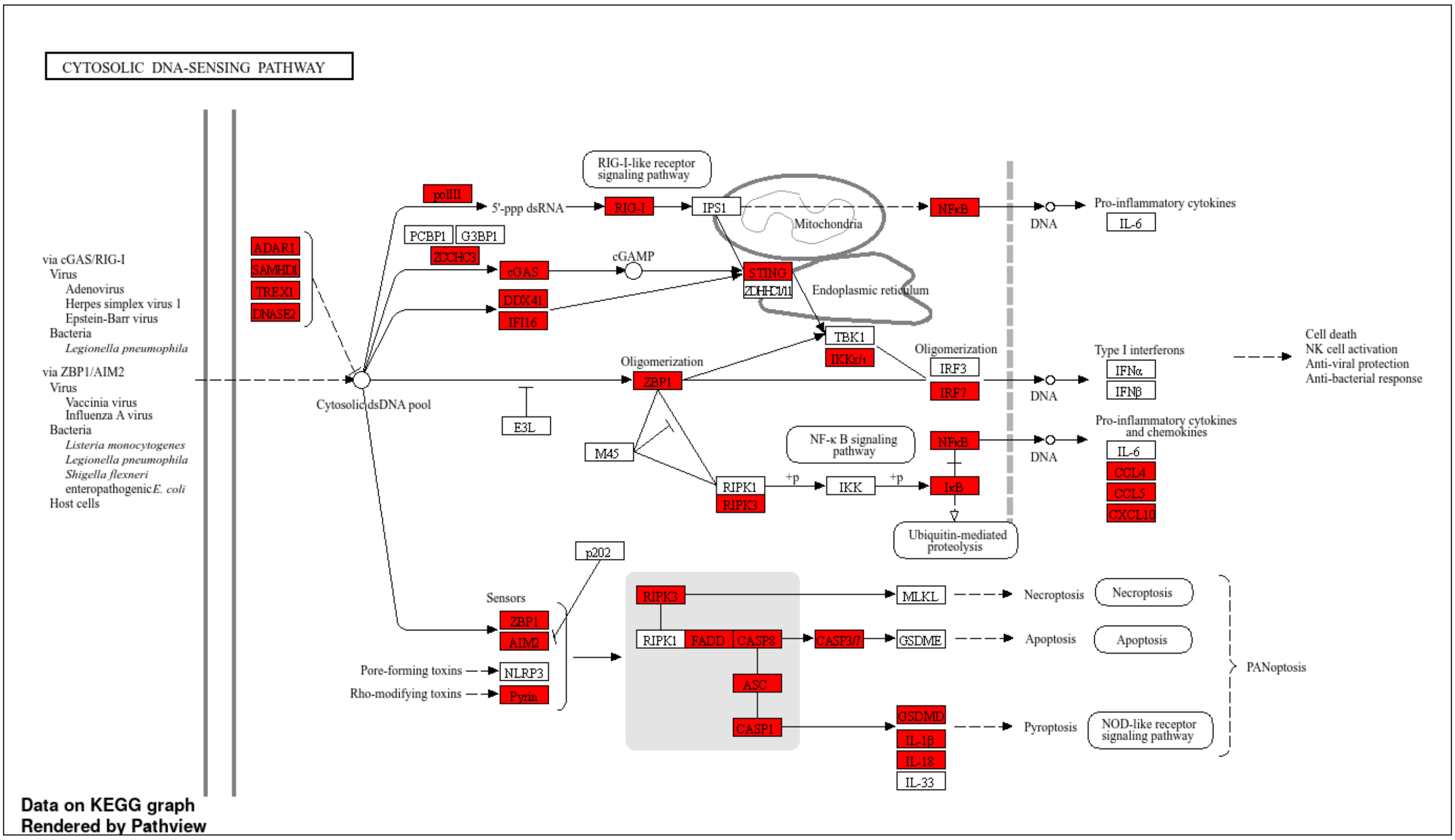

### Suppl. Figure S3

A

B

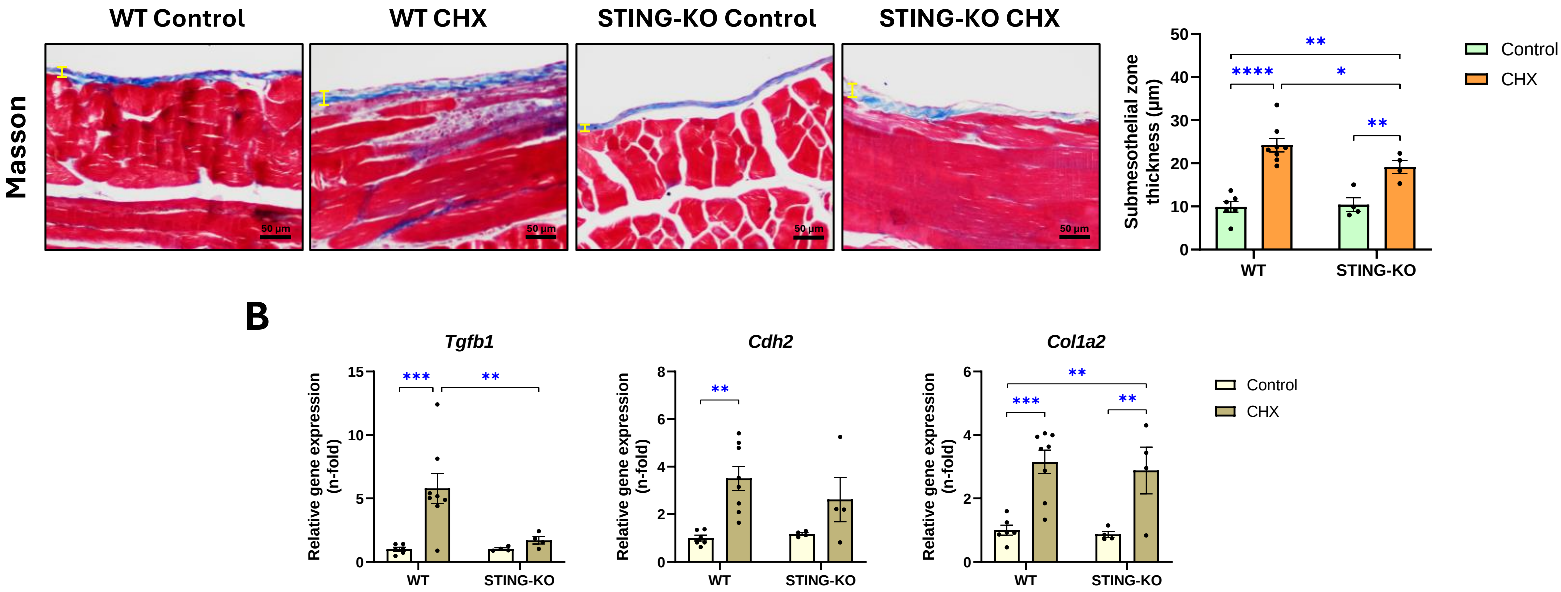

### Suppl. Figure S4

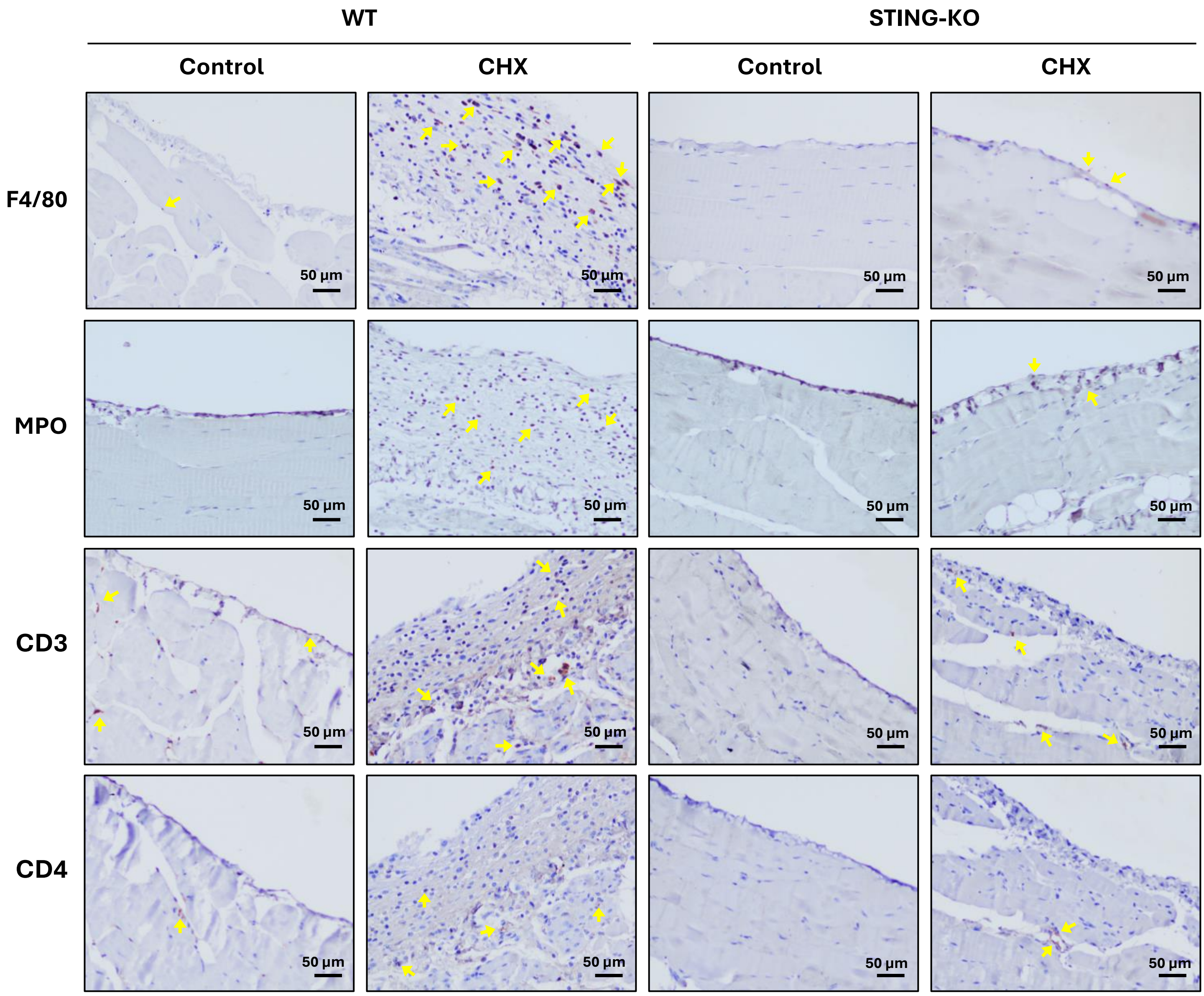

### Suppl. Figure S5

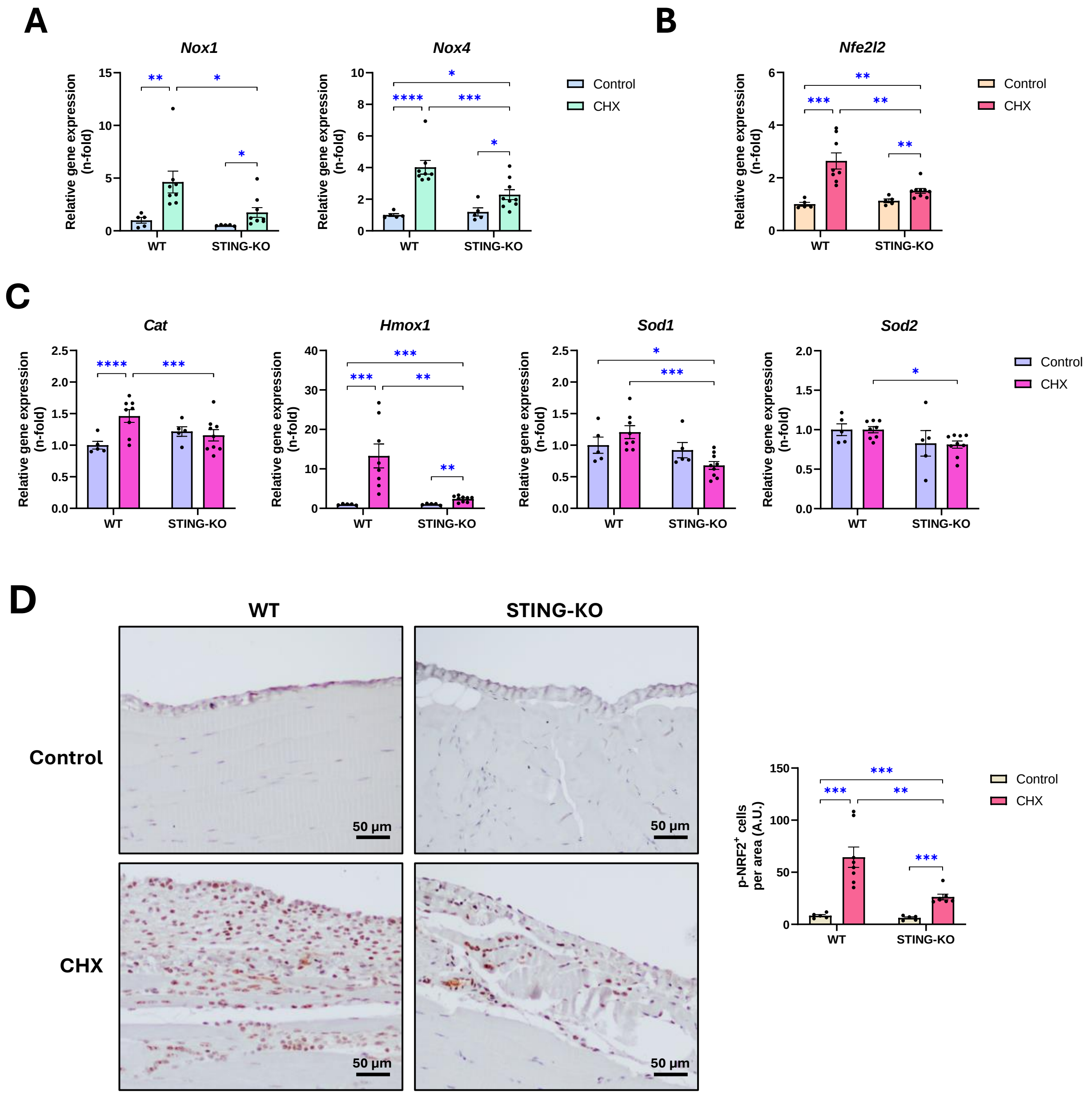

### Suppl. Figure S6

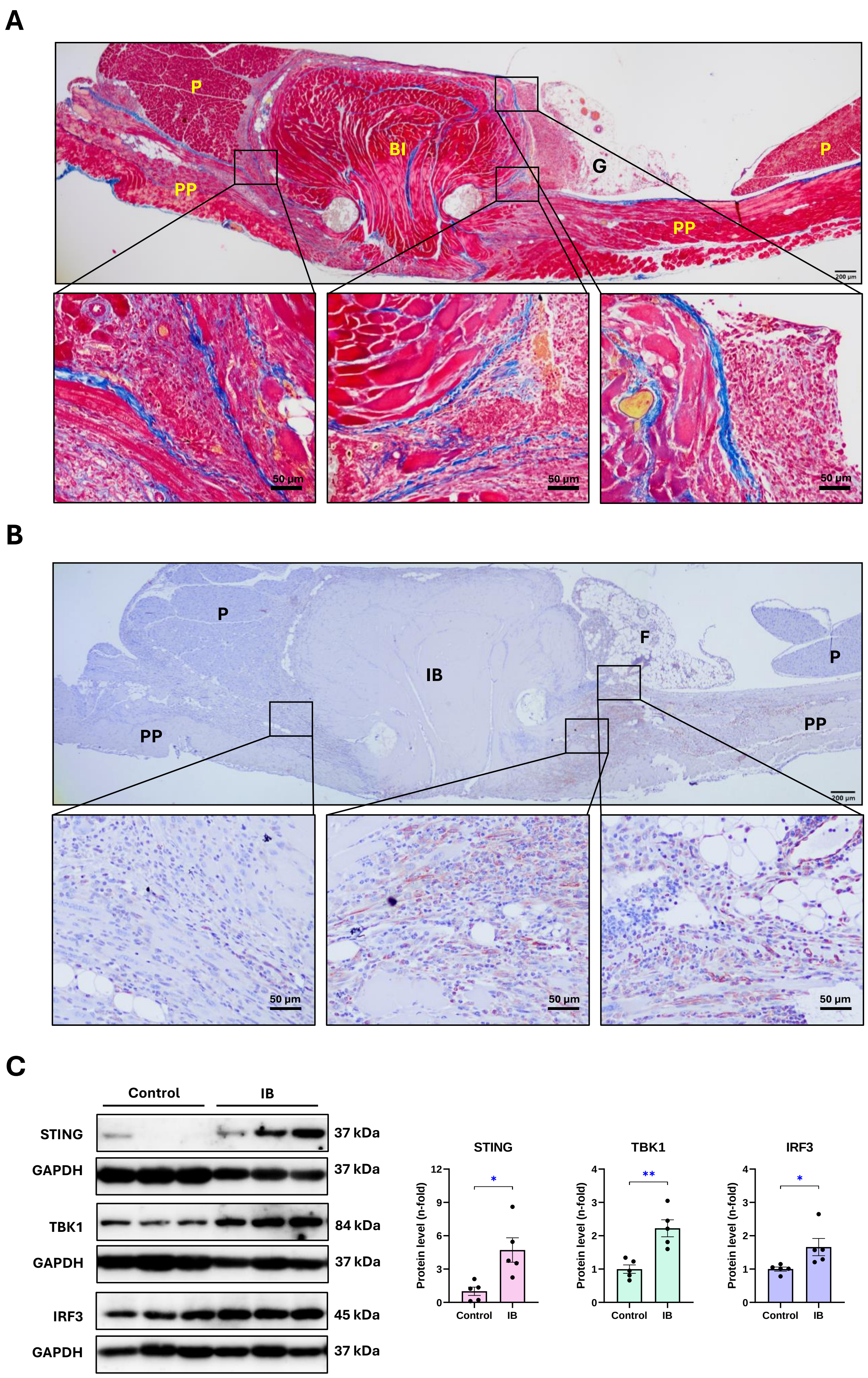

### Suppl. Figure S7

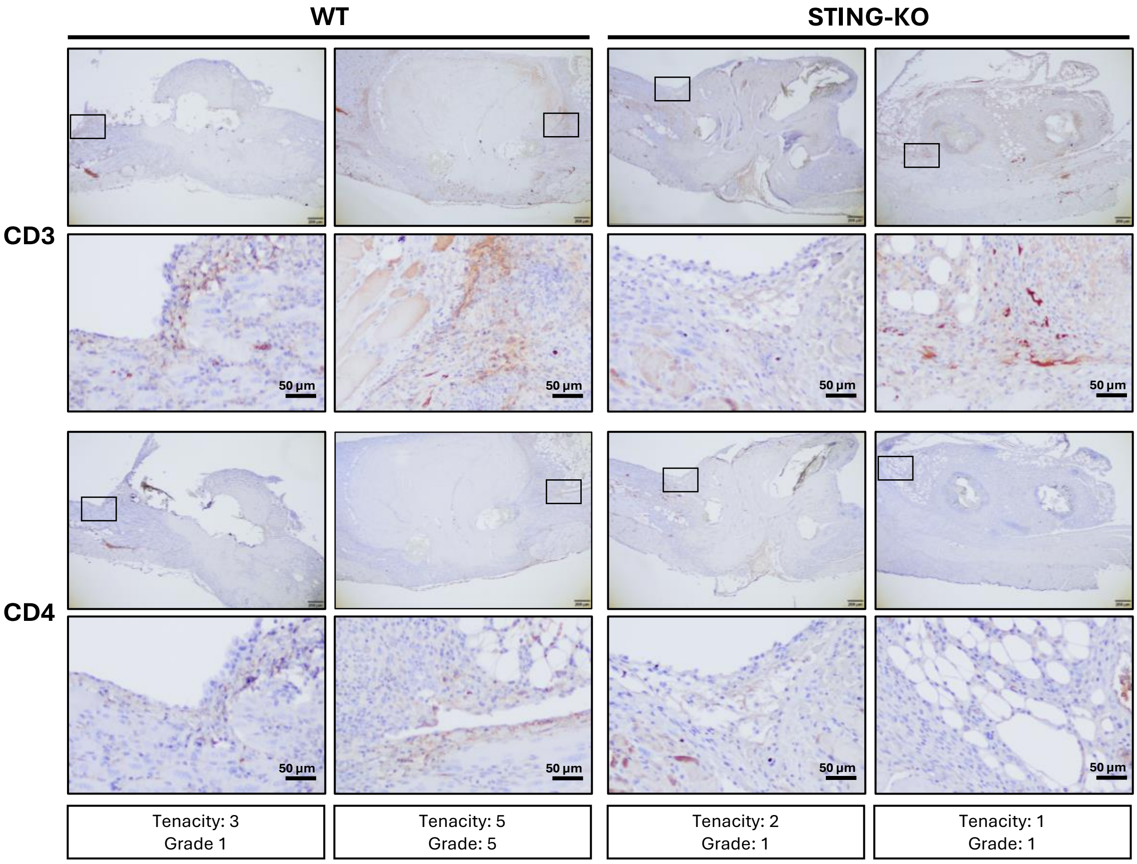

#### Supplementary Figure Legends

**Supplementary Figure S1. Time course of the peritoneal damage induced by CHX in mice.** Male C57BL/6J mice were daily intraperitoneally injected with 0.1% CHX for 3, 10, or 30 days (d). **(A)** Peritoneal membrane thickness assessment. The figure shows Masson's trichrome stained parietal peritoneal tissue sections of representative mice from each group (left) and the corresponding quantification of submesothelial zone thickness (right). The yellow lines indicate the width measured. Scale bar: 50 µm. **(B,C)** Relative gene expression of mesothelial-to-mesenchymal transition (MMT) and fibrosis markers **(B)** and proinflammatory chemokines and cytokines **(C)** analyzed by RT-qPCR from total RNA of parietal peritoneal tissue, using *Gapdh* as housekeeping gene, and expressed as fold change (n-fold) relative to control. Results are represented as mean ± SEM of 6 animals per group. * $p < 0.05$; ** $p < 0.01$; *** $p < 0.001$; and **** $p < 0.0001$ versus control (Ctrl). + $p < 0.05$; ++ $p < 0.01$; +++ $p < 0.001$; and ++++ $p < 0.0001$ versus CHX 3 d; # $p < 0.05$; and ### $p < 0.001$ versus CHX 10 d.

**Supplementary Figure S2. Upregulated genes of the cytosolic DNA-sensing pathway in the peritoneum of 10-day-CHX-exposed mice.** The figure shows the mmu04623: Cytosolic DNA-sensing pathway - Mus musculus generated by the Kyoto Encyclopedia of Genes and Genomes (KEGG) database. Genes of this pathway that were significantly upregulated (q-value < 0.05) in the transcriptomic analysis performed in the parietal peritoneal tissue of mice treated with 0.1% chlorhexidine gluconate (CHX) for 10 days are highlighted in red.

**Supplementary Figure S3. Peritoneal membrane thickness and fibrosis in WT and STING-KO exposed to CHX for 10 days.** Male C57BL/6J wild-type (WT) and STING-deficient (STING-KO) mice were intraperitoneally injected with 0.1% chlorhexidine gluconate (CHX) for 10 days. **(A)** Peritoneal membrane thickness assessment. The figure shows Masson's trichrome stained parietal peritoneal tissue sections of representative mice from each group (left) and the corresponding quantification of the submesothelial zone thickness (right). The yellow lines indicate the width measured. Scale bar: 50 µm

**(B)** Relative gene expression of mesothelial-to-mesenchymal transition (MMT) and fibrosis markers analyzed by RT-qPCR from total RNA of parietal peritoneal tissue, using *Gapdh* as housekeeping gene. Results are expressed as n-fold relative to control and are represented as mean ± SEM of 4-8 animals per group. * $p < 0.05$; ** $p < 0.01$; *** $p < 0.001$; and **** $p < 0.0001$.

**Supplementary Figure S4. Inflammatory cell infiltration into the peritoneum of WT and STING-KO mice exposed to CHX for 30 days.** Male C57BL/6J wild-type (WT) and STING-deficient (STING-KO) mice were intraperitoneally injected with 0.1% chlorhexidine gluconate (CHX) for 30 days. Parietal peritoneal tissue sections were used for immunohistochemistry staining to identify inflammatory immune cell subsets using specific antibodies against the following markers: F4/80 (macrophages), myeloperoxidase (MPO, neutrophils), and CD3 and CD4 (lymphocytes). Microscopy images correspond to a representative animal from each group. Cells with positive staining are pointed out by arrows. Scale bar: 50 µm.

**Supplementary Figure S5. Peritoneal oxidative stress and antioxidant response in WT and STING-KO mice exposed to CHX for 30 days.** Male C57BL/6J wild-type (WT) and STING-deficient (STING-KO) mice were intraperitoneally injected with 0.1% chlorhexidine gluconate (CHX) for 30 days. **(A-C)** Relative gene expression levels of the oxidative enzymes NAPDH oxidases 1 and 4 (*Nox1* and *Nox4*) **(A),** the transcription factor NRF2 (*Nfe2l2* gene) **(B)**, and the antioxidant response enzymes catalase (*Cat*), heme oxygenase 1 (*Hmox1*), and superoxide dismutases (*Sod1* and *Sod2*) **(C).** Relative gene expression levels were analyzed by RT-qPCR from total RNA of parietal peritoneal tissue, using *Gapdh* as a housekeeping gene, and expressed as fold change (n-fold) relative to the control group. **(D)** Immunohistochemical staining of phosphorylated NRF2 (p-NRF2) in parietal peritoneal tissue sections. Microscopy images correspond to a representative animal from each group. Scale bar: 50 µm. Results are represented as the mean ± SEM of 5–9 animals per group. * $p < 0.05$; ** $p < 0.01$; *** $p < 0.001$; and **** $p < 0.0001$. A.U.: arbitrary units.

**Supplementary Figure S6. Expression of STING and downstream signaling mediators in postsurgical peritoneal adhesions.** Ischemic buttons (IBs) were surgically generated in the peritoneum of male C57BL/6J mice and the formation of adhesions between peritoneal IBs and surrounding tissues was assessed after 5 days from the surgery. **(A)** Masson's trichrome staining in sections of parietal peritoneal tissue containing IBs. Peritoneal adhesions of the parietal peritoneum (PP) with neighbor organ and tissues, like pancreas (P) and fat (F) are shown. Squared areas containing peritoneal adhesion interfaces are shown at a higher magnitude below. **(B)** Immunohistochemical detection of STING in sections of parietal peritoneal tissue containing IBs. The figure shows $STING^+$ cells of the peritoneal peritoneum based near peritoneal adhesions. Squared areas containing $STING^+$ cells are depicted at a higher magnitude below. Microscopy images correspond to an IB with adhesions with grade and tenacity scores equal to 5. Scale bars: 200 and 50 µm. **(C)** Protein levels of STING, TBK1, and IRF3 assessed by western blot from total protein of parietal peritoneal tissues, using GAPDH as loading control. Results are represented as fold change (n-fold) relative to the control group and expressed as mean ± SEM of 5 animals per group. * $p < 0.05$ and ** $p < 0.01$.

**Supplementary Figure S7. T cell infiltration in peritoneal adhesions of WT and STING-KO mice.** Three ischemic buttons (IBs) were surgically made in the peritoneum of male C57BL/6J wild-type (WT) and STING-deficient (STING-KO) mice. The formation of adhesions on the IBs was assessed 5 days after surgery. Sections of parietal peritoneal tissue containing the IBs were used for immunohistochemical detection of $CD3^+$ or $CD4^+$ T cells. Microscopy images correspond to two representative animals from each group. The squared areas are shown at higher magnification below every original micrograph. Scale bars: 200 and 50 µm.
